# Piezo1 agonist restores meningeal lymphatic vessels, drainage, and brain-CSF perfusion in craniosynostosis and aged mice

**DOI:** 10.1101/2023.09.27.559761

**Authors:** Matt J. Matrongolo, Phillip S. Ang, Junbing Wu, Aditya Jain, Josh K. Thackray, Akash Reddy, Chi Chang Sung, Gaëtan Barbet, Young-Kwon Hong, Max A. Tischfield

## Abstract

Skull development coincides with the onset of cerebrospinal fluid (CSF) circulation, brain-CSF perfusion, and meningeal lymphangiogenesis, processes essential for brain waste clearance. How these processes are affected by craniofacial disorders such as craniosynostosis are poorly understood. We report that raised intracranial pressure and diminished CSF flow in craniosynostosis mouse models associates with pathological changes to meningeal lymphatic vessels that affect their sprouting, expansion, and long-term maintenance. We also show that craniosynostosis affects CSF circulatory pathways and perfusion into the brain. Further, craniosynostosis exacerbates amyloid pathology and plaque buildup in *Twist1^+/−^:5xFAD* transgenic Alzheimer’s disease models. Treating craniosynostosis mice with Yoda1, a small molecule agonist for Piezo1, reduces intracranial pressure and improves CSF flow, in addition to restoring meningeal lymphangiogenesis, drainage to the deep cervical lymph nodes, and brain-CSF perfusion. Leveraging these findings, we show Yoda1 treatments in aged mice with reduced CSF flow and turnover improve lymphatic networks, drainage, and brain-CSF perfusion. Our results suggest CSF provides mechanical force to facilitate meningeal lymphatic growth and maintenance. Additionally, applying Yoda1 agonist in conditions with raised intracranial pressure and/or diminished CSF flow, as seen in craniosynostosis or with ageing, is a possible therapeutic option to help restore meningeal lymphatic networks and brain-CSF perfusion.

## Introduction

Meningeal lymphatic vessels (MLVs) facilitate the drainage of cerebrospinal fluid (CSF) from the head, and in doing so help control central nervous system (CNS) waste clearance, immune surveillance, and responses to injury (1). MLVs reside in dura mater where they grow along the venous sinuses in close apposition to the skull. They express many transcription factors and cell surface markers common to lymphatic vessels. However, the growth of MLVs occurs exclusively during the first few postnatal weeks in mice, and in a basal to dorsal progression from the base to the top of the skull, mirroring the establishment of CSF circulation routes (2–4). VEGF-C/VEGFR3 signaling is required for the growth of MLV networks but unlike most peripheral lymphatic vessels, MLVs require continuous VEGF-C signaling in order to maintain vessel integrity and survival (2). MLVs are further distinguished from peripheral lymphatics according to transcriptional regulation (5). These data suggest the growth and maintenance of MLVs are dictated by processes unique to the specialized meningeal environment where these vessels reside. Yet, the environmental factors that shape how these vessels develop and are maintained in dura are still poorly understood.

The growth and expansion of lymphatic networks is dependent upon increasing interstitial fluid and laminar flow (6–8). Laminar flow also helps maintain the integrity of mature lymphatic networks via activation of Piezo1-mediated mechanotransduction signaling (7). MLVs that grow in perisinusoidal dura alongside the venous sinuses have access to CSF and thus exposure to laminar flow (9–11), although mechanisms that channel CSF and its contents through the arachnoid membrane are not well understood. Nonetheless, injury models that impede CSF drainage to the deep cervical lymph nodes (dCLNs) are associated with pathological changes to MLVs, potentially because these vessels try to compensate for loss of flow (12, 13). As CSF flow and turnover naturally declines with age (14, 15), dorsal MLVs along the superior sagittal sinus (SSS) and confluence regress, whereas vessels found at the skull base along the petrosquamousal sinus become hyperplastic and take on a lymphedematous-like appearance (16, 17). These observations suggest CSF may help guide meningeal lymphangiogenesis and subsequently the maintenance of mature networks to facilitate CNS waste clearance and immune surveillance.

Restoring the functional integrity of MLVs in pathological conditions or with ageing has important clinical implications. In aged animals, rejuvenation of MLVs with adenoviral delivery of VEGF-C improves brain-CSF perfusion, the drainage of macromolecules from CSF, and normal cognitive functions (17). In traumatic brain injury models, this application also limits the extent of Iba1 gliosis (12). In mouse Alzheimer’s disease (AD) models, augmenting MLV functions with VEGF-C enhances the effectiveness of monoclonal antibodies that target amyloid-beta to clear plaques from the brain (18). In intracranial tumor models, improving MLV drainage with VEGF-C promotes survival and generates immunogenic responses against tumors (19, 20). Thus, stimulating MLV expansion and facilitating drainage to the dCLNs can enhance waste clearance and immune surveillance with ageing and in disease states, underscoring the need to further characterize factors that control the growth, maturation, and maintenance of these vessels.

Given that MLVs develop in periosteal dura attached to the skull, we previously investigated their growth and expansion in *Twist1^FLX/FLX^:Sm22a-Cre* mouse models for craniosynostosis (CS), a prevalent craniofacial disorder that affects skull growth via premature fusion of the cranial sutures (21, 22). In humans, loss-of-function mutations in the transcription factor *TWIST1* or gain-of-function mutations in *FGFR2* cause syndromic forms of CS that are associated with dural venous sinus malformations and raised intracranial pressure (ICP) (23–25). We showed that the growth and sprouting of dorsal MLVs along the transverse sinuses was reduced in *Twist1^FLX/FLX^:Sm22a-Cre* mice, whereas basal vessels were hyperplastic in some animals, and drainage to the dCLNs was diminished (22). Insults to the surrounding dural extracellular matrix and/or loss of growth factor signaling from venous smooth muscle was postulated to affect MLV sprouting and expansion.

In the present study, we show that pathological changes to MLVs across multiple CS models occur without affection to dura or the venous sinuses, and instead are associated with raised ICP and diminished CSF flow to perisinusoidal dura. In addition to reduced MLV sprouting and expansion, adult mice with CS show premature regression of dorsal MLVs and hyperplastic basal vessels, suggesting MLVs are exposed to conditions in CS that induce precocious ageing. Reduced drainage of CSF macromolecules to the dCLNs in CS mice associates with reduced CSF perfusion into the brain. Additionally, crossing CS mice with 5xFAD models for familial AD causes increased amyloid-beta buildup in the brain. We show that treating CS mice with Yoda1, a selective Piezo1 agonist that reduces the mechanical threshold for channel activation, reduces ICP and helps restore the growth and sprouting of MLVs, drainage to the dCLNs, and brain-CSF perfusion. Finally, activating Piezo1 with Yoda1 in aged mice, during which time CSF flow and turnover is reduced, also improves MLV coverage, drainage, and brain-CSF perfusion. Our results suggest that raised ICP in CS restricts MLV access to CSF and affects their development, maintenance, and functional drainage, while also hindering brain-CSF perfusion and macromolecule clearance. Thus, MLVs appear sensitive to changes in laminar flow, suggesting CSF may provide mechanical force to facilitate the development, integration, and maintenance of CNS waste clearance systems across the lifespan.

## Results

### Intracranial pressure is increased in mouse models for Twist1 CS

Raised ICP is a primary concern in CS, causing neurological complications and cognitive deficits if left untreated (24, 25). Injury models have shown that raised ICP is associated with transient, pathological changes to MLV morphology and affects drainage of CSF macromolecules to the deep cervical lymph nodes (dCLNs) (12, 13). However, the effects of pathological, chronically raised ICP–as seen in conditions such as CS–on MLV development, their long-term maintenance, and drainage to the dCLNs has not been investigated. We addressed this question by conditionally inactivating a single copy of *Twist1* using *Sm22a-Cre*, both of which are expressed in periosteal dura and sutural mesenchyme, but not MLVs (22, 23). Approximately 56% (18/32) of *Twist1^+/FLX^:Sm22a-Cre* adult mice had unilateral or bilateral coronal synostosis, with partial or full fusion of the sutures, the former more common (**Figure 1A-C**). We next measured ICP and found it was significantly increased, similar to previous findings in *Twist1^+/−^* mice, which are also used to model syndromic CS (26) (**Figure 1F**). In agreement, ICP was also increased in our *Twist1^+/−^* mouse models (**Figure 1F**). Coronal synostosis was also more penetrant in adult *Twist1^+/−^* mice (∼76%, 23/30) with a greater proportion showing full unilateral and/or bilateral synostoses (**Figure 1D**). In *Twist1^+/−^* mice without suture fusion, ICP was normal (**Figure 1E, F**). Thus, both *Twist1^+/FLX^:Sm22a-Cre* and *Twist1^+/−^* mice have suture fusion that associates with raised ICP, as found in human CS.

**Figure 1:**
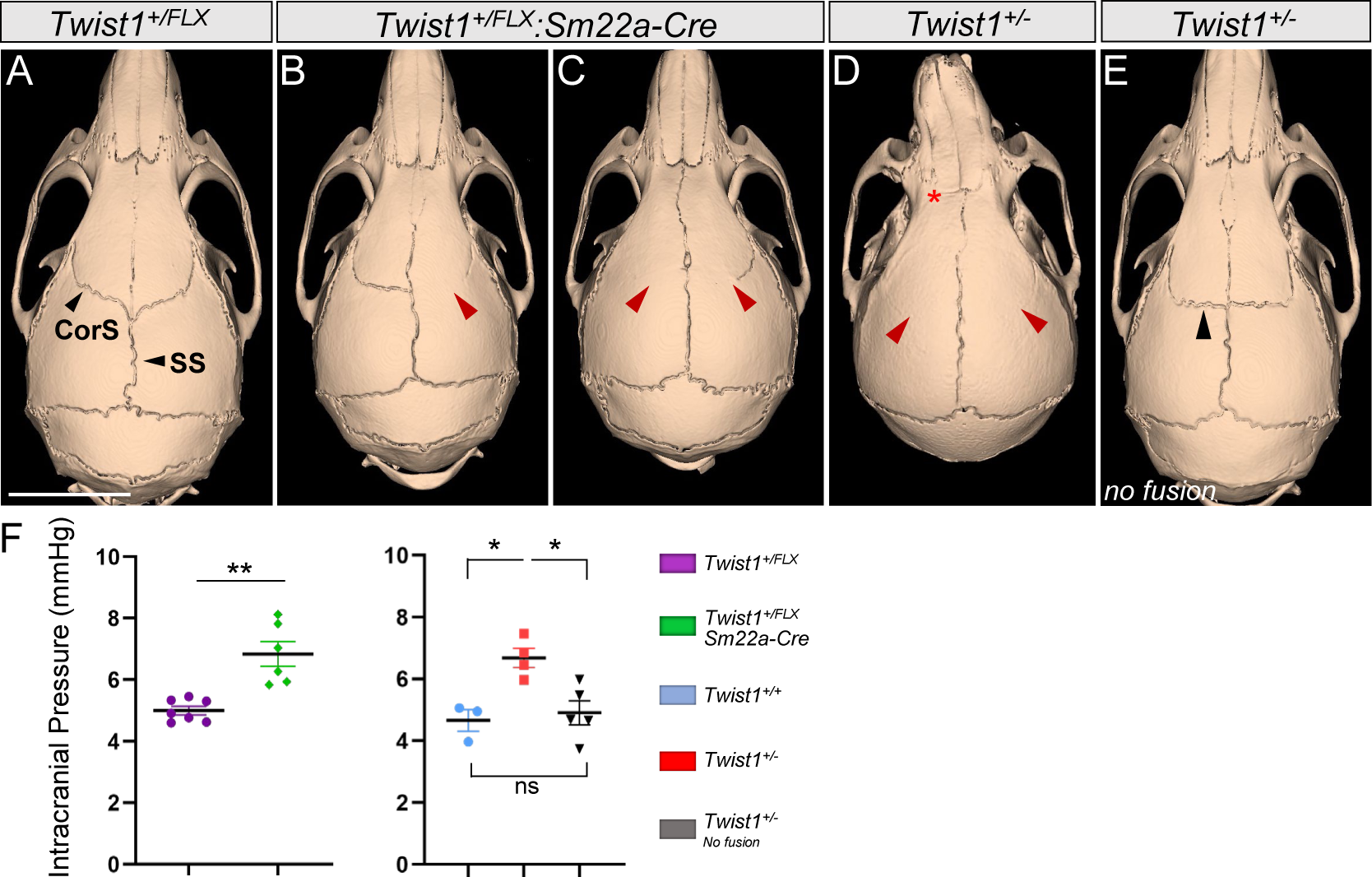
Twist1 haploinsufficiency in cranial sutural mesenchyme causes CS and raised ICP. Representative reconstructed computed tomography (CT) scans from two-month-old adults showing normal skull and suture morphology in a *Twist1^+/FLX^* control (A) versus *Twist1^+/FLX^:Sm22a-Cre* (B and C) and *Twist1^+/−^* mice with (D) or without (E) suture fusion. (B) Near-complete unilateral fusion of the right coronal suture (red arrowhead). (C) Partial bilateral fusion with full and partial fusion on the left and right coronal sutures, respectively (red arrowheads). (D) Complete bilateral fusion (red arrowheads) accompanied by fusion of the frontonasal suture (asterisk) and a deviated nasal bone, more common in *Twist1^+/−^* mice. (E) *Twist1^+/−^* mouse without suture fusion. In the absence of fusion, the orientation of the coronal sutures is flatter and more ‘box-like’ (black arrowhead). (F) Intracranial pressure is raised in *Twist1^+/FLX^:Sm22a-Cre* and *Twist1^+/−^*mice with CS but is normal in *Twist1^+/−^* mice without suture fusion [*Twist1^+/FLX^ (n=7); Twist1^+/FLX^:Sm22a-Cre (n=6); Twist1^+/+^ (n=3); Twist1^+/−^ (n=4); Twist1^+/−^ no fusion (n=5)*]. CorS=coronal suture; SS=sagittal suture. \**p≤0.05,* \*\**p≤0.01.* two-tailed unpaired t test with Welch’s correction (left) one way ANOVA with Tukey’s multiple comparison test (right). Scale bar=5mm.

### CS causes MLV growth and sprouting deficits in the absence of dura and venous sinus malformations

We previously reported that the sprouting and expansion of dorsal MLVs along the transverse sinuses was reduced in homozygous *Twist1^FLX/FLX^:Sm22a-Cre* mice. These mice are more severely affected than heterozygous *Twist1^+/FLX^:Sm22a-Cre* and *Twist1^+/−^* mice, with regionalized absence of mineralized bone in the calvarium, hypoplastic dura, and loss/hypoplasia of the transverse sinuses (22). We speculated that hypoplasia of the dural extracellular matrix and/or loss of growth factor signaling from venous smooth muscle may have exerted deleterious effects on MLVs.

The transverse sinuses were not missing nor hypoplastic in *Twist1^+/FLX^:Sm22a-Cre* and *Twist1^+/−^* mice and showed normal smooth muscle coverage, the latter of which is also consistent with *Twist1^FLX/FLX^:Sm22a-Cre* homozygotes **(Figure S1A)** (22). The dura and arachnoid membranes, as labeled by Crabp2, also appeared qualitatively normal at the dorsal midline in newborn *Twist1^+/−^* pups **(Figure S1B)**. In juveniles and adults, the dural membrane was not hypoplastic and could be peeled off the skull in one piece **(Figure S1C)**, which was impossible in *Twist1^FLX/FLX^:Sm22a-Cre* homozygotes due to its hypoplastic state. Given the absence of overt changes to dura and the surrounding venous vasculature as seen in *Twist1^FLX/FLX^:Sm22a-Cre* mice, we asked whether MLV growth and sprouting was affected in *Twist1^+/FLX^:Sm22a-Cre* and *Twist1^+/−^* animals at postnatal day 17 (P17), at which time MLVs are actively growing and sprouting in dura.

Numerous loops and sprouts were present in juvenile control mice along the transverse sinuses, especially at or near regions where ‘hotspots’ are found in adults (**Figure 2A**). Hotspots are important for immune cell and macromolecule uptake and show increased branching and complexity along the transverse sinuses and at the confluence (5). Interestingly, the growth and sprouting of MLV networks was affected in both *Twist1^+/FLX^:Sm22a-Cre* and *Twist1^+/−^* mice, similar to homozygous *Twist1^FLX/FLX^:Sm22a-Cre* mice. MLV coverage and average vessel diameter was reduced along the transverse sinuses, as was growth at the sinus confluence and superior sagittal sinus. Sprouting along the transverse sinuses was significantly reduced, especially at locations where hotspots were forming (**Figure 2A, A’**). Similar to adults, ICP was already increased by this time (**Figure 2A’**). Thus, CS perturbs the growth and sprouting of MLVs, even with the absence of dura and venous sinus malformations.

**Figure 2:**
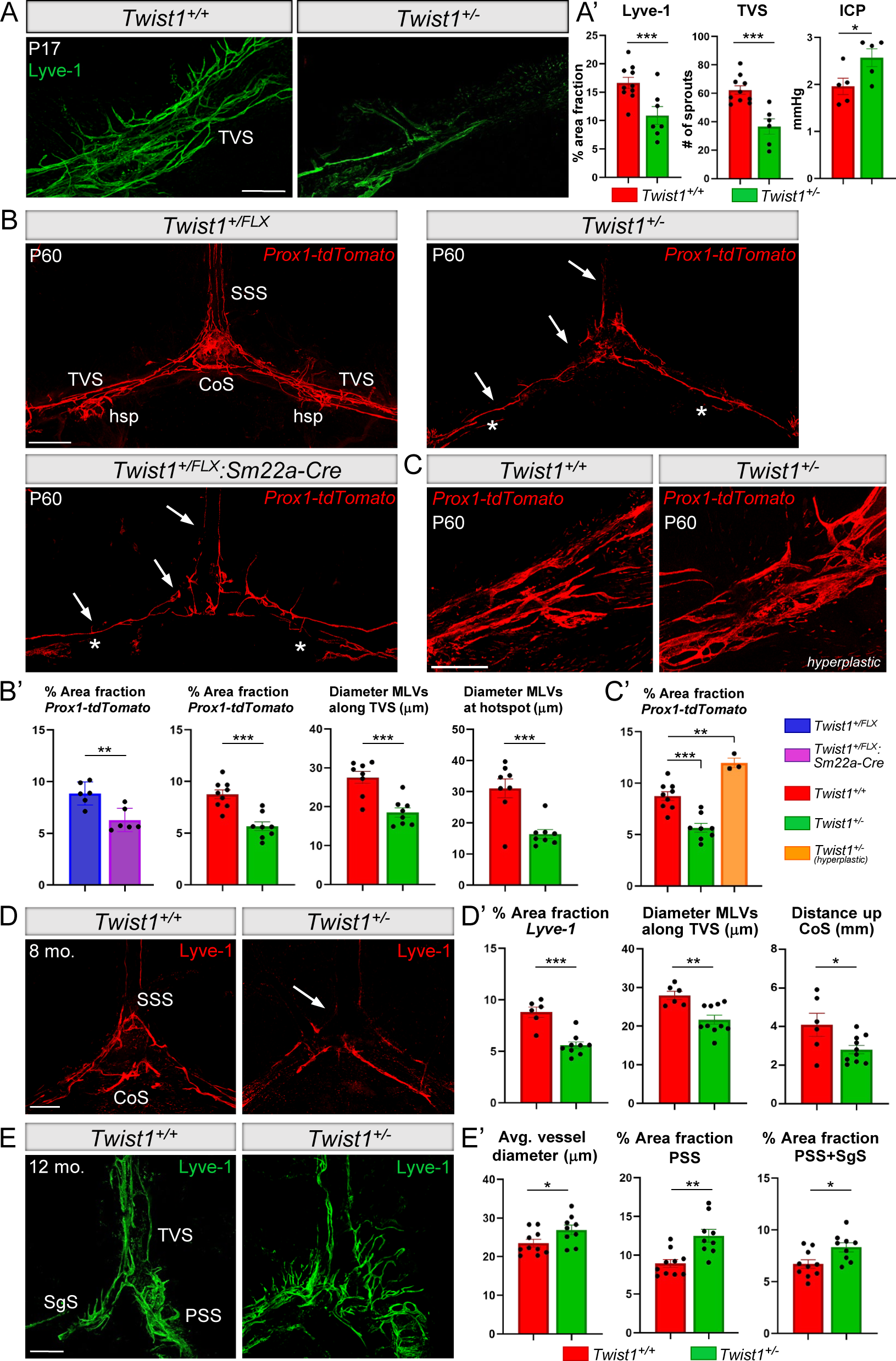
CS affects MLV sprouting, expansion, and long-term maintenance. (A) Representative images from juvenile P17 mice. Sprouting and expansion of dorsal MLVs along the TVS is reduced in mice with CS and raised ICP. (A’) Quantification of percent area fraction of Lyve-1 [*Twist1^+/+^* (n=10); *Twist1^+/−^* (n=7)], number of sprouts [*Twist1^+/+^* (n=10); *Twist1^+/−^* (n=6)], and levels of ICP at P17 [n=5/genotype]. (B) Compared with controls (top left), dorsal MLVs along the SSS and TVS (arrows) are hypoplastic in *Twist1^+/FLX^:Sm22a-Cre:Prox1-tdTomato* (bottom left) and *Twist1^+/−^ :Prox1-tdTomato* mice (top right) with CS. Hotspots along the TVS are poorly formed or completely missing, as shown in these mice with bilateral absence (asterisks). (B’) Quantification of percent area fraction [*Twist1^FLX/+^* and *Twist1^FLX/+^:Sm22a-Cre* (n=6); *Twist1^+/+^* (n=9) and *Twist1^+/−^* (n=8)] and average vessel diameters along the TVS [*Twist1^+/+^* and *Twist1^+/−^* (n=8)] and at hotspots [*Twist1^+/+^* and *Twist1^+/−^* (n=8)]. (C and C’) A small subset of CS mice show signs of vessel hyperplasia [*Twist1^+/−^* (n=3)]. (D) In 8-10-month-old adults, dorsal MLVs are further regressed, especially at the sinus confluence and along the SSS (arrow). (D’) Quantification of Lyve-1 percent area fraction [*Twist1^+/+^* (n=6); *Twist1^+/−^* (n=9)], average vessel diameter along the TVS [*Twist1^+/+^* (n=6); *Twist1^+/−^* (n=10)], and growth from the CoS [*Twist1^+/+^* (n=6); *Twist1^+/−^* (n=10)]. (E) Basal MLV networks along the SgS and PSS are hyperplastic in *Twist1^+/−^*mice with CS. (E’) Quantification of average vessel diameter and the percent area fraction of MLVs along the PSS and SgS [*Twist1^+/+^* (n=10); *Twist1^+/−^* (n=9)]. SSS=superior sagittal sinus; TVS=transverse sinus; CoS=confluence of sinuses; hsp=hotspot; SgS=sigmoid sinus; PSS=petrosquamosal sinus. \**p≤0.05*, \*\**p≤0.01, ***p≤0.001* two-tailed unpaired t test (B’, D’, E’), one-way ANOVA with Tukey’s multiple comparison test (C’). Scale bar: (A) 200μm; (B) 1mm; (C, D, E) 500μm

### MLVs show morphological changes in young adult mice with CS and raised ICP

Our previous findings in two-three-month-old *Twist1^FLX/FLX^:Sm22a-Cre* adults revealed hypoplastic dorsal MLV networks along the transverse and superior sagittal sinuses (22). In two-four-month-old *Twist1^+/FLX^:Sm22a-Cre* and *Twist1^+/−^*adults, dorsal networks along the transverse and superior sagittal sinuses were also hypoplastic with reduced branching and complexity (**Figure 2B, B’, Figure S1E)**. Compared with *Twist1^FLX/FLX^:Sm22a-Cre* mice, findings were similar but typically less severe. Hotspots along the transverse sinuses were either hypoplastic or missing on one or both sides (**Figure 2B**). Vessel complexity and area coverage at the sinus confluence and along the superior sagittal sinus were also reduced. These findings were more common in mice that had bilateral coronal synostosis or full unilateral coronal synostosis. By contrast, dorsal networks were usually less affected in more mildly affected mice with partial unilateral fusion and, notably, MLVs were normal in mice without synostosis and raised ICP (**Figure S2A, B**). Interestingly, MLVs along the pterygopalatine and middle meningeal arteries appeared to develop normally, even in animals with full bilateral fusion, despite poor development of neighboring vessels patterned along the venous sinuses **(Figure S1D)**. These results suggest MLVs along arterial vessels are spared in CS, and that their growth, expansion, and/or maintenance might be differentially regulated from those that grow along the venous sinuses.

To determine if MLVs show pathological changes in different genetic forms of CS, we developed a model for Apert Syndrome using mice that permit conditional activation of a constitutively active Fgfr2 receptor (*Fgfr2^S252W^*) that is the most common mutation found in syndromic CS (27). *Fgfr2^+/S252W^:Sm22a-Cre* adult mice had full to partial fusion of the sagittal suture and full to partial, unilateral or bilateral, fusion of the coronal sutures **(Figure S3A)**. ICP was significantly increased in adults **(Figure S3A’)**. Furthermore, dorsal MLV networks were hypoplastic and also missing hotspots **(Figure S3B, B’)**. Thus, pathological changes to MLVs are conserved across other forms of syndromic CS with raised ICP.

### Dorsal MLV networks prematurely regress as CS mice age

The majority of two-four-month-old adult mice with full unilateral or partial/full bilateral fusion had hypoplastic dorsal networks. On rare occasions, however, some mice showed signs of vessel hyperplasia at the sinus confluence and/or at hotspots along the transverse sinus (**Figure 2C, C’**). Interestingly, following blunt trauma to the head, acutely raised ICP can increase vessel coverage, the numbers of loops and sprouts, and vessel diameter at or near hotspots (12). These phenotypes revert back to baseline approximately one month later, by which time ICP has normalized. This suggests dorsal MLV networks are undergoing certain pathological changes that may be influenced by the type/degree of suture fusion and/or variations in the levels of ICP. In addition, the levels of ICP increase as *Twist1^+/−^* adult mice age (26), suggesting pathological changes to MLVs may continue to progress in CS. We therefore characterized the effects of chronically raised ICP on dorsal MLV networks in middle-aged *Twist1^+/−^* mice at eight-to-ten-months. Compared with younger adults, dorsal MLV networks were more affected. Hotspots along the transverse sinus were rudimentary or completely missing and vessel coverage at the sinus confluence and proximally along the superior sagittal sinus was further reduced (**Figure 2D, D’**). Remarkably, these phenotypes are similar to mice aged ∼20-24 months (16, 17), although they are present at least one year earlier in CS models. Thus, dorsal MLVs continue to regress and ‘age’ prematurely in CS.

### Basal MLVs demonstrate early onset hyperplasia in CS

MLVs near the skull base along the petrosquamousal and sigmoid sinuses are a major outflow route for CSF macromolecules to the dCLNs **(Figure S1E)** (16). Although dorsal networks show regression in aged mice, basal lymphatics become hyperplastic for reasons that are still unclear. We previously reported that basal networks were hyperplastic in a subset of *Twist1^FLX/FLX^:Sm22a-Cre* mice (22). We analyzed basal MLVs in *Twist1^+/−^* animals using a preparation that preserves these vessels in whole mounts (22). In two-four-month-old adult mice, basal networks appeared normal, despite affection to dorsal networks. However, basal networks were hyperplastic in middle-aged *Twist1^+/−^* mice (8-12 months), especially along the petrosquamosal sinus (**Figure 2D, D’**). This result is striking because wild-type mice only show this phenotype by ∼24 months of age (16), at which time CSF flow and turnover is reduced and drainage to the dCLNs declines (14, 17). Thus, MLVs in middle-aged mice with CS show signs of precocious ageing, with premature regression of dorsal vessels coupled with hyperplastic basal networks.

### CSF flow to perisinusoidal dura and the dCLNs is reduced in CS

The growth and sprouting of peripheral lymphatics require increasing ISF and laminar flow (7, 8). Thus, we hypothesized CS and raised ICP may affect the flow of CSF to perisinusoidal dura, potentially impinging upon the development of MLVs along the venous sinuses, which receive access to CSF. We injected a 45kDa ovalbumin-647 conjugated tracer into the cisterna magna of *Twist1^+/FLX^:Sm22a-Cre* and *Twist1^+/−^* mice at P17 and sacrificed them one hour later. We detected tracer accumulation at locations where hotspots are typically located (**Figure 3A**). This suggests hotspots are forming in juvenile mice before weaning. In *Twist1^+/FLX^:Sm22a-Cre* and *Twist1^+/−^* mice, diminished growth and sprouting of MLVs was associated with reduced flow to the perisinusoidal dura (**Figure 3A, C**). In two-to-four-month-old controls, tracer was also localized to branched hotspots along the transverse sinuses and at the sinus confluence (**Figure 3B**). In CS animals, we again found significantly less tracer in perisinusoidal dura surrounding dorsal and basal MLVs and less accumulation at hotspots (**Figure 3C, Figure S2B, see also Figure S1E for orientation)**. These results suggest limiting CSF access to perisinusoidal dura affects the growth and sprouting of MLVs in CS.

**Figure 3:**
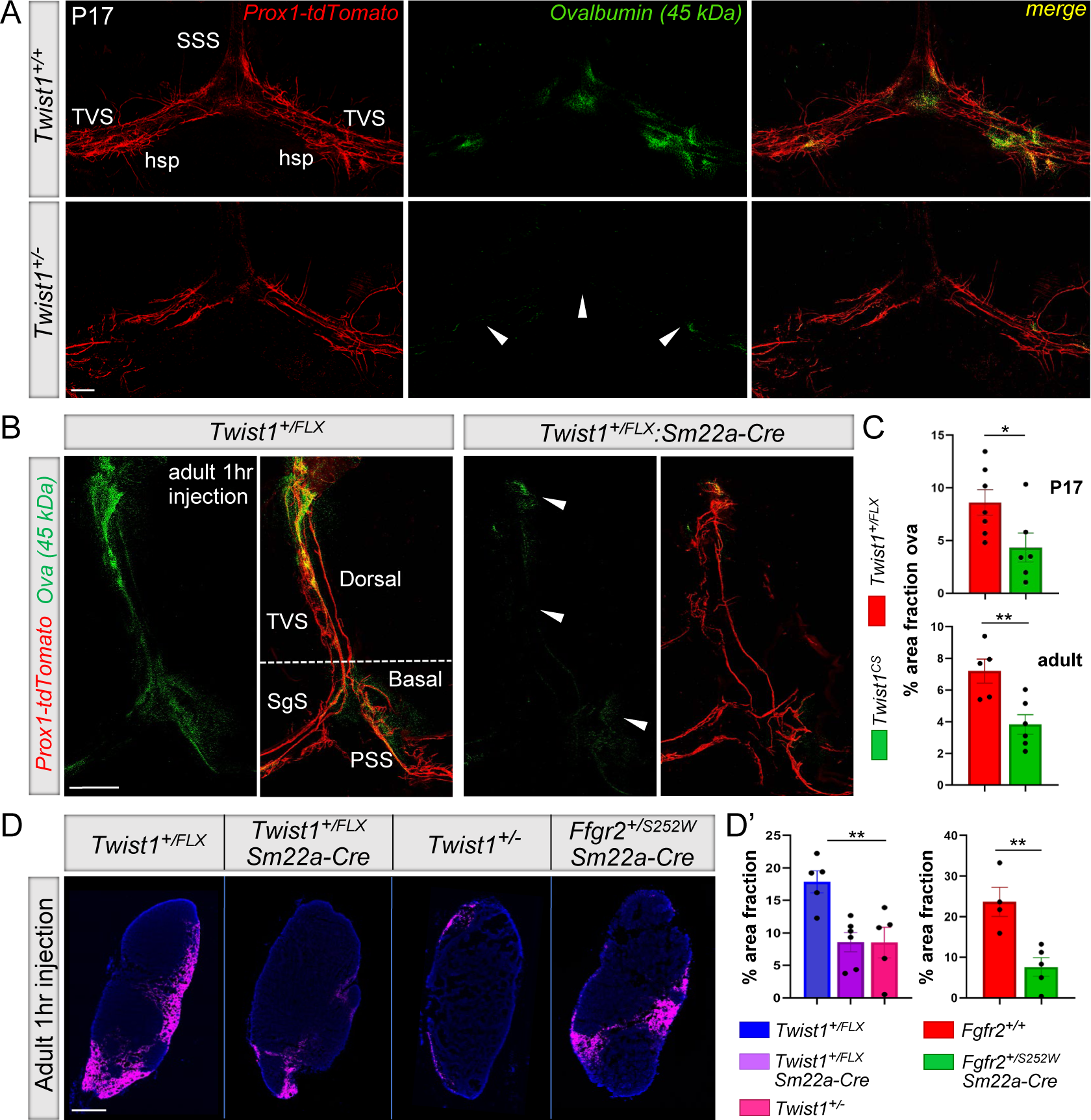
CSF flow to perisinusoidal dura and the dCLNs is reduced in mice with CS. (A and C) Following injection of 45kDa ovalbumin tracer into the cisterna magna, juvenile P17 mice with *Twist1* haploinsufficiency and CS (bottom panel in A) show a significant reduction in tracer deposition in perisinusoidal dura compared to unaffected littermates (top panel), especially at regions where hotspots form (arrowheads) [*Twist1^+/+^* (n=7); *Twist1^+/−^* (n=6)]. (B and C) In 2-4-month-old adults, tracer deposition in perisinusoidal dura surrounding dorsal MLVs is significantly reduced. Flow to perisinusoidal dura surrounding basal vessels is also reduced (arrowheads) [*Twist1^FLX/+^* (n=5); *Twist1^FLX/+^:Sm22a-Cre* (n=6)]. See figure S1E for additional orientation. (D) Representative images show that tracer drainage to the dCLNs is significantly reduced in *Twist1^+/FLX^:Sm22a-Cre*, *Twist1^+/−^*, and *Fgfr2^+/S252W^:Sm22a-Cre* mice with CS. (D’) Quantification of percent area fraction of ovalbumin tracer in dCLNs [*Twist1^FLX/+^* (n=5); *Twist1^FLX/+^:Sm22a-Cre* (n=6); *Twist1^+/−^* (n=5); *Fgfr2^+/+^* (n=4); *Fgfr2^+/S252W^:Sm22a-Cre* (n=5)]. SSS=superior sagittal sinus; TVS=transverse sinus; hsp=hotspot; SgS=sigmoid sinus; PSS=petrosquamosal sinus. \**p≤0.05*, \*\**p≤0.01* two-tailed unpaired t test (C, D’ left). One-way ANOVA with Tukey’s multiple comparison test (D’ right). Scale bar: (A) 500μm; (B) 200μm; (D) 200μm

We previously showed that MLV deficits in adult *Twist1^FLX/FLX^:Sm22a-Cre* homozygous mice were associated with less drainage of CSF macromolecules to the dCLNs (22). We asked whether drainage to the dCLNs was similarly reduced in two-four-month-old *Twist1^+/FLX^:Sm22a-Cre*, *Twist1^+/−^*, and *Fgfr2^+/S252W^:Sm22a-Cre* mice. We injected a 45kDa ovalbumin tracer into the cisterna magna and sacrificed the mice one hour later. The dCLNs showed less tracer in CS mice versus unaffected controls (**Figure 3D**). Tracer accumulation in the dCLNs and also perisinusoidal dura was normal in *Twist1^+/−^* mice without suture fusion, consistent with normal levels of ICP and intact MLV networks **(Figure S2A’-D’)**. Thus, drainage of macromolecules in CSF to the dCLNs is impaired across different genetic models for CS.

These results prompted us to reassess our original findings in homozygous *Twist1^FLX/FLX^:Sm22a-Cre* mice (22). Like *Twist1^+/FLX^:Sm22a-Cre* and *Twist1^+/−^* mice, ICP was significantly increased and there was less tracer in perisinusoidal dura **(Figure S4A, B)**. This indicates that pathological changes to MLVs associate with reduced access to CSF across different CS models. Overall, these changes were generally more severe in *Twist1^FLX/FLX^:Sm22a-Cre* mice compared to what we observed in *Twist1^+/FLX^:Sm22a-Cre* or *Twist1^+/−^* mice, presumably due to the additive effects that insults from the surrounding environment exerted on top of flow deficits.

### MLVs can access CSF via flow through the cerebellar tentorium

Following tracer injection into the cisterna magna of control mice, we perfused animals with fixative and hemi-dissected the skull along the dorsal midline to visualize outflow routes. As previously reported, tracer flow was detected at the cribriform plate and also along the skull base where MLVs line the dural sheaths of exiting cranial and spinal nerves (28–30). Tracer was also concentrated around MLVs at the rostral rhinal venous plexus. Interestingly, comet-like streaks of tracer were evident in the cerebellar tentorium and these appeared to terminate at dorsal and basal hotspots (**Figure 4A, A’**). We did not see tracer deposition in dura where MLVs grow alongside the middle meningeal arteries, consistent with previous reports (5) (**Figure S1D**). This is notable considering arterial MLVs are not affected in young adult CS mice versus those vessels that grow along the venous sinuses and have access to CSF. Through careful preservation of the cerebellar tentorium (**Figure 4C**), we observed that lymphatic vessels traversed regions where CSF tracer accumulates, and these vessels appeared to terminate at lymphatic hotspots located along the transverse sinuses (**Figure 4B**). Some tracer was found inside these vessels, whereas most was engulfed by dural macrophages, as previously reported (5) (**Figure 4B, B’**). We also examined uptake by injecting a fluorescently conjugated Lyve-1 antibody into the CSF via the cisterna magna. Antibody labeling was detected in basal and dorsal vessels (especially at hotspots), and some labeling was also apparent in vessels traversing the tentorium one-hour post-injection (**Figure 4B’, B”**). These findings are intriguing because the mechanisms by which MLVs access CSF are unclear, and it has been speculated that a subset of MLVs may contact the arachnoid and/or there may be regionalized breaks in the barrier that allow CSF to exit (31). Perhaps more consistent with the latter, MLVs can receive access to CSF via flow through the cerebellar tentorium.

**Figure 4:**
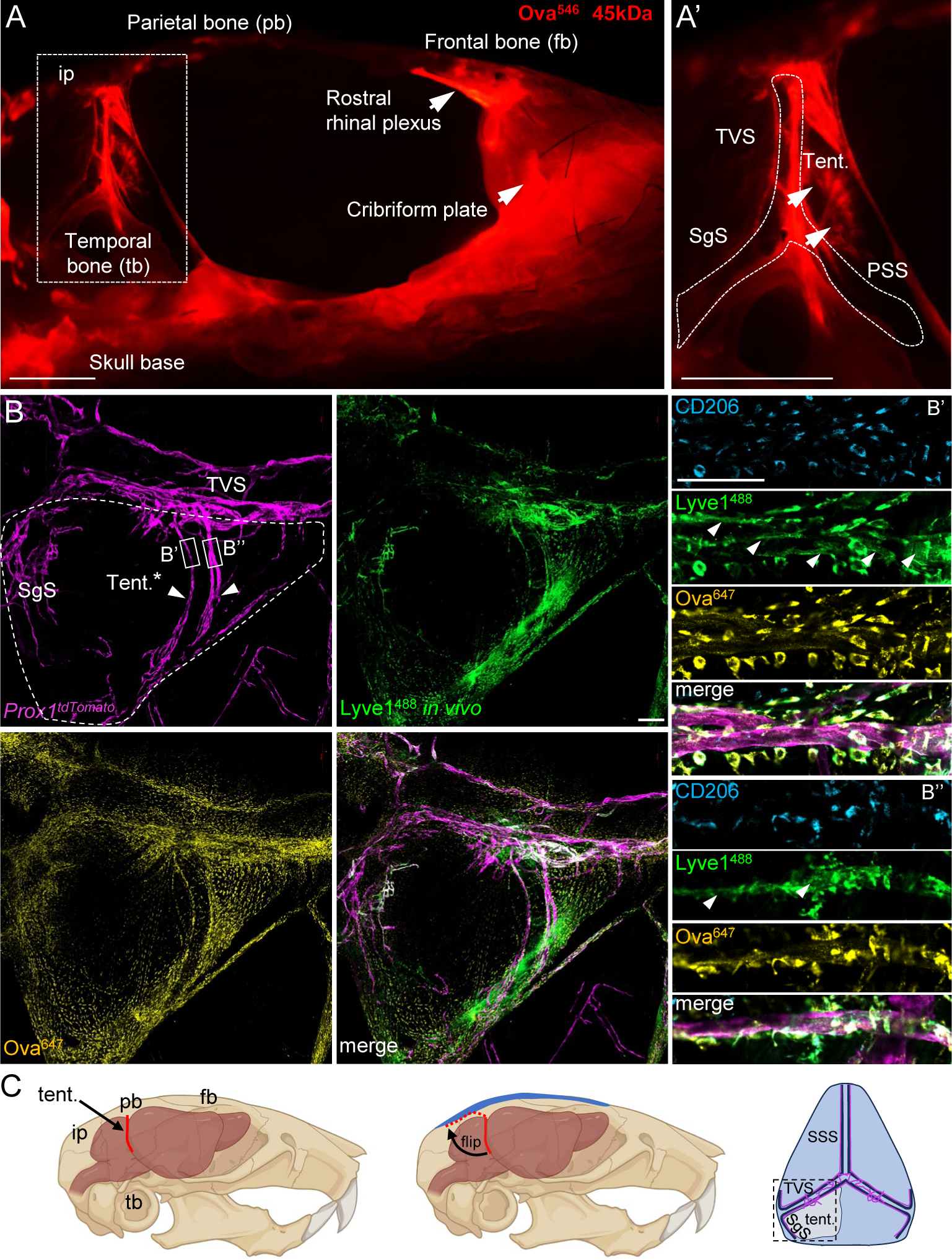
MLVs can access CSF via flow through the cerebellar tentorium. (A) Adult skull hemi-dissected along the dorsal midline. Following injection of 45kDa ovalbumin tracer into the cisterna magna, tracer is seen draining through the cribriform plate and the skull base. Comet-like streaks of tracer are also seen in the cerebellar tentorium (tent.), as demarcated by arrowheads in the magnified boxed image (A’). (B) Compressed z-stacks of a dural flatmount showing MLVs labeled with *Prox1-tdTomato* traversing the tentorium (denoted by arrowheads). These vessels form extensions with lymphatic hotspots located along the transverse sinus (TVS). Injecting conjugated Lyve-1^488^ antibody (green) or 45kDa ovalbumin tracer (gold) into CSF via the cisterna magna shows labeling in the tentorium and also MLVs located at hotspots along the TVS and sigmoid sinus (SgS). (B’ and B”) Corresponding magnified images of boxed regions of interest in panel B. MLVs traversing the tentorium show staining for Lyve-1^488^ (in vivo, green, arrowheads) and 45kDa ovalbumin. Neighboring CD206+ dural macrophages are also labeled by the antibody and engulf the tracer. (C) Schematic overviews illustrating the orientation of the tentorium in panel B. The tentorium, a dural inflection that separates the cortex from the cerebellum and brainstem, was carefully preserved, flipped backwards during coverslipping, and flat mounted on dura that underlies the interparietal bone (ip) for visualization purposes (dotted red outline). Pb=parietal bone; Fb=frontal bone; Tb=temporal bone; SSS=superior sagittal sinus. Scale bar: (A, A’) 2.5mm; (B) 200μm; (B’) 100μm

### CS affects CSF circulation and influx of CSF macromolecules into the brain

Genetic, pharmacological, and surgical methods that ablate MLVs or block drainage to the dCLNs diminish paravascular (i.e., glymphatic) influx of CSF macromolecules into the brain (17). Loss of CSF tracer in perisinusoidal dura in CS mice prompted us to investigate if CSF circulation and CSF-brain perfusion were affected. We co-injected 3kDa dextran and 45kDA ovalbumin tracers into the cisterna magna and visualized tracer flow in CSF by transcranial live imaging through the skull (32). In unaffected controls, tracer preferentially flowed along previously reported pathways (4, 33). Tracer was first detected along dorsal routes, with accumulation seen in the pineal recess and in the tentorium where MLVs reside. Approximately 15-20 minutes later, tracer in basal circulation pathways began to pool in the olfactory recess. Around this time, tracer was also seen in the paravascular spaces surrounding penetrating pial arteries, indicative of brain perfusion by CSF (34) (**Figure 5A**).

**Figure 5:**
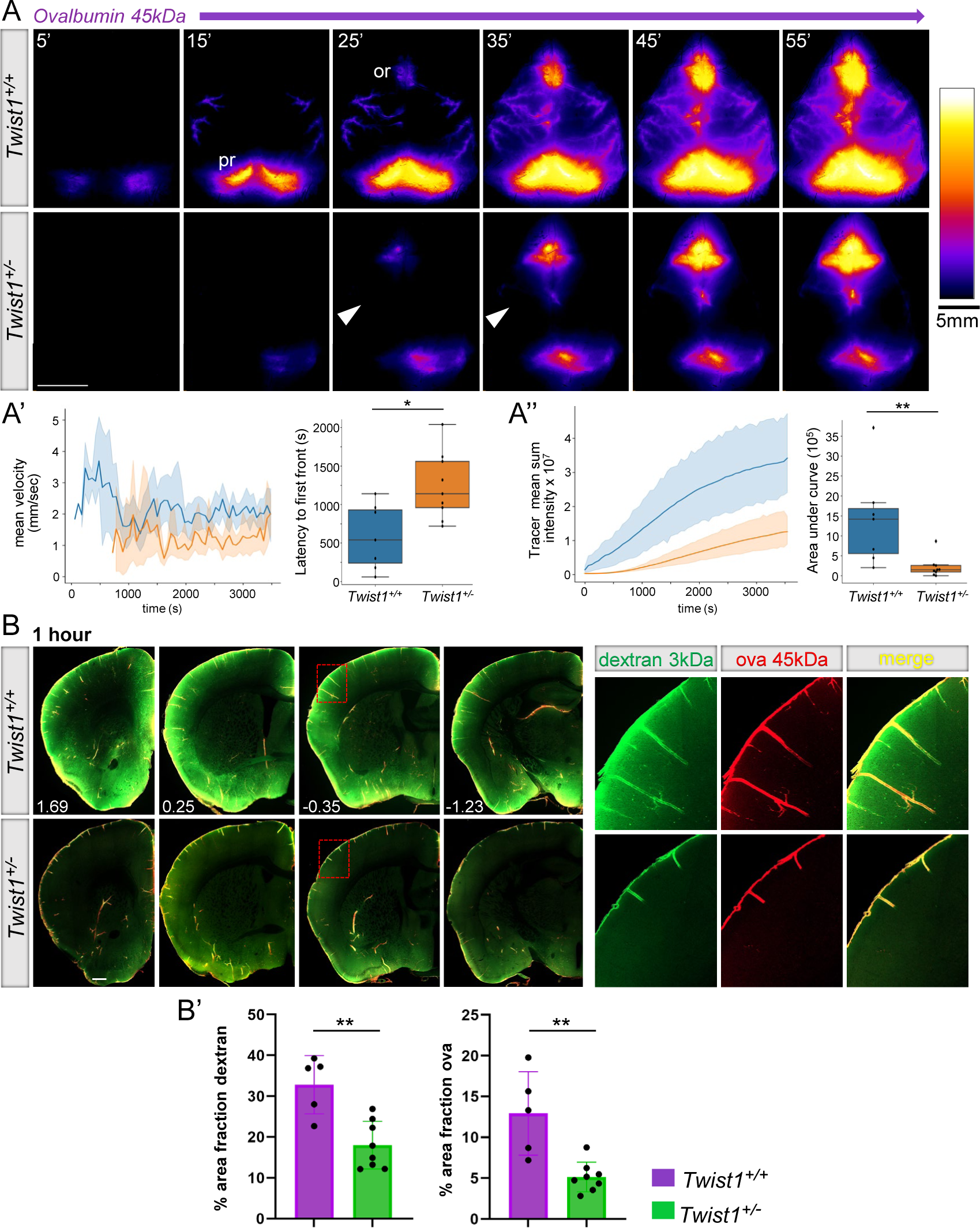
CS affects CSF flow and the perfusion of CSF macromolecules into the brain. (A) Representative images taken during transcranial live imaging of a 45kDa ovalbumin tracer injected into the cisterna magna of young adult mice. Compared to unaffected littermates (top panels), *Twist1^+/−^* mice with CS (bottom panels) show a delay in the appearance of CSF tracer along preferred dorsal pathways (A’). Throughout one-hour imaging sessions, the mean sum intensity of tracer was reduced (A’’), especially in the pineal recess and paravascular spaces surrounding penetrating pial arteries (arrowheads) [*Twist1^+/+^* (n=7); *Twist1^+/−^* (n=9)]. (B) Perfusion of CSF macromolecules into the brain is reduced in *Twist1^+/−^* mice. Magnified boxed images on the right show that tracer labeling along penetrating arteries is shallower in *Twist1^+/−^*mice compared to unaffected littermates. (B’) Quantification of percent area fraction of 3kDa Dextran and 45kDa Ovalbumin in brain slices [*Twist1^+/+^* (n=5); *Twist1^+/−^* (n=8)]. \**p≤0.01,* \*\**p≤0.01* (A’, A’’) Mann-Whitney U test with Bonferroni correction. (B’) two-tailed unpaired t test. Scale bar: (A) 5mm; (B) 500μm

Given that suture fusion was more penetrant in *Twist1^+/−^* versus *Twist1^+/FLX^:Sm22a-Cre* mice, we focused our efforts on the former. In adult *Twist1^+/−^* mice, there was typically a five-to-ten-minute delay in the appearance of tracer flow along dorsal routes, with less signal seen in the pineal recess and along the dorsal cortex (**Figure 5A-A’’**). Basal flow was less affected in some mice as judged by pooling in the olfactory recess. Labeling along penetrating pial arteries, especially along more dorsal regions of the cortex, was also diminished (**Figure 5A**). Thus, CS appears to perturb CSF circulation along preferred dorsal pathways.

Postmortem examination of brain tissue from *Twist1^+/−^* CS mice showed a significant reduction of both the 3kDa dextran and 45kDA ovalbumin tracers (**Figure 5B, B’**). The depth of tracer labeling along penetrating pial arterioles was shallower compared to unaffected littermates. Similar findings were also seen in postmortem brains from *Fgfr2^+/S252W^:Sm22a-Cre* animals **(Figure S5A, B)**. By contrast, CSF-brain perfusion was unaffected in *Twist1^+/−^* mice without suture fusion, suggesting deficits in flow and CSF-brain perfusion are linked to synostosis **(Figure S2E, E’)**. These results suggest CS impedes CSF-brain perfusion and the clearance of waste and macromolecules from the brain.

### AQP4 polarization to glial endfeet is reduced along penetrating cortical vessels in CS

Paravascular influx of CSF and glymphatic waste exchange is dependent upon AQP4 water channels, which are polarized to glial endfeet that form the glial limitans (34). Injecting dextran and ovalbumin tracers of various sizes into CSF of AQP4 knockout mice results in a significant loss of influx and penetration beneath the brain surface (35). We therefore asked whether the polarization of AQP4 to glial endfeet was affected in CS. We first looked at the cortex of P17 CS mice, at which time AQP4 polarization to glial endfeet is nearing completion (4). We found a subtle, albeit significant, reduction along large caliber vessels **(Figure S6A)**. Biochemical fractionation of blood vessels with tethered glial endfeet from whole brain tissue also revealed a reduction in AQP4 protein **(Figure S6B, B’)**. In adult animals with CS, however, we did not detect obvious differences in AQP4 by immunolabeling, suggesting AQP4 polarization to glial endfeet was delayed at earlier time points. Nonetheless, co-labeling with lectin and AQP4 still showed a significant reduction along larger caliber vessels **(Figure S6C, C’)**. Thus, CS appears to delay and/or affect the polarization of AQP4 to glial endfeet abutting large cortical vessels, which may exert subtle effects on brain-CSF perfusion. However, this does not appear to be a major factor affecting perfusion in our models, and instead changes to preferred CSF circulation pathways and/or insults to MLVs and drainage to the dCLNs seem more likely (17).

### CS exacerbates amyloid-beta pathology in 5xFAD mice

Genetic, surgical, and pharmacological models that inhibit MLV drainage impair brain-CSF perfusion and the clearance of macromolecules from the brain (13, 17, 28). Also, light-activated ablation of MLVs using Visudyne aggravates amyloid-beta pathology in *5xFAD* mice, which express five human mutations found in Alzheimer’s disease (AD) and develop amyloid-beta plaques (17). We investigated whether loss of MLV drainage and brain-CSF perfusion in CS exacerbated amyloid-beta deposition by crossing *Twist1^+/−^* and *5xFAD* mice. In five-six-month-old *Twist1^+/−^:5xFAD* mice, there was a significant increase in area coverage, number of amyloid-beta plaques, and plaque size in both the cortex and hippocampus compared to *5xFAD* mice without CS (**Figure 6A, B**). Immunostaining for GFAP and IBA1 in brain sections from *Twist1^+/−^* CS mice revealed normal GFAP and IBA1 coverage, and also microglia morphology, suggesting no changes to astro- and microgliosis under steady state **(Figure S7)**. Notably, suture fusion was very mild or non-existent in *Twist1^+/FLX^:Sm22a-Cre:5xFAD* mice, presumably due to background strain differences. We also did not see a significant increase in area coverage, number of amyloid-beta plaques, and aggregate size in either the cortex or hippocampus (**Figure. 6C, C’**). This suggests cranial suture fusion and corresponding deficits in brain-CSF perfusion and meningeal lymphatic drainage are linked to amyloid-beta buildup in *Twist1^+/−^:5xFAD* mice with CS.

**Figure 6:**
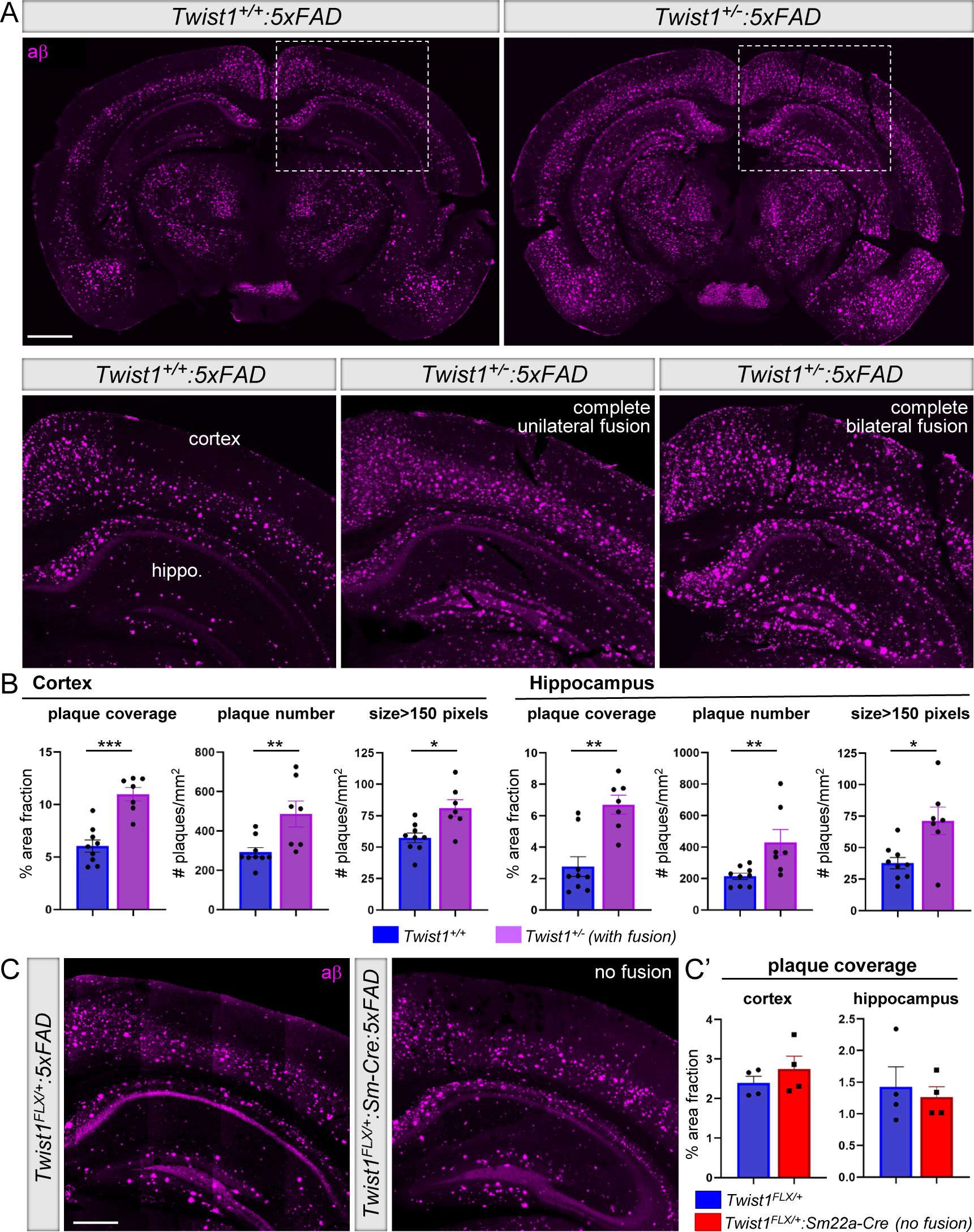
CS exacerbates amyloid-β pathology in 5xFAD transgenic mice. (A) Representative images of amyloid-β in the cortex and hippocampus of 5-6-month-old 5xFAD mice with CS (top right and bottom panels) and without CS (top left). (B) Quantification of amyloid plaque coverage, number, and size in the cortex and hippocampus [*Twist1^+/+^:5xFAD* (n=9); *Twist1^+/^:5xFAD^−^*(n=7)]. (C) Representative images of amyloid-β in the cortex and hippocampus of 5-month-old *5xFAD* and *Twist1^FLX/+^:Sm22a-Cre:5xFAD* mice without suture fusion. (C’) Quantification of percent area fraction of amyloid-β in the cortex and hippocampus. No differences are seen in amyloid-β coverage [*Twist1^FLX/+^* and *Twist1^FLX/+^:Sm22a-Cre* (n=4)]. \**p≤0.05*, \*\**p≤0.01, ***p≤0.001* Mann-Whitney U-test. Scale bar: (A) 1mm; (C) 500 μm

### Piezo1 activation reduces ICP and helps improve brain-CSF perfusion and meningeal lymphatic functions in CS mice

Raised ICP in CS is postulated to result from several factors, including craniocerebral disproportion and/or venous hypertension coupled with changes to cerebral blood flow (25). Piezo1, a mechanosensitive ion channel expressed in vascular endothelial cells and smooth muscle, is suggested to control cerebral capillary blood flow, vascular tone, and blood pressure (36, 37), implying it may be involved in the regulation of ICP. Recent studies have also indicated a role for Piezo1 in the regulation of intraocular pressure (38). In addition, force exerted by unidirectional laminar flow activates Piezo1-mediated mechanotransduction signaling in lymphatic vessels to control sprouting, expansion, and long-term maintenance of lymphatic networks (7). Given that we could also detect Piezo1 expression in meningeal blood vessels and perisinusoidal MLVs using *Piezo1^tdTomato^* reporter mice (**Figure. 7A, B**), we tested whether activating Piezo1 with a small molecule agonist could reduce the levels of ICP and/or help restore brain-CSF perfusion and meningeal lymphatic functions in *Twist1^+/−^* CS mice.

**Figure 7:**
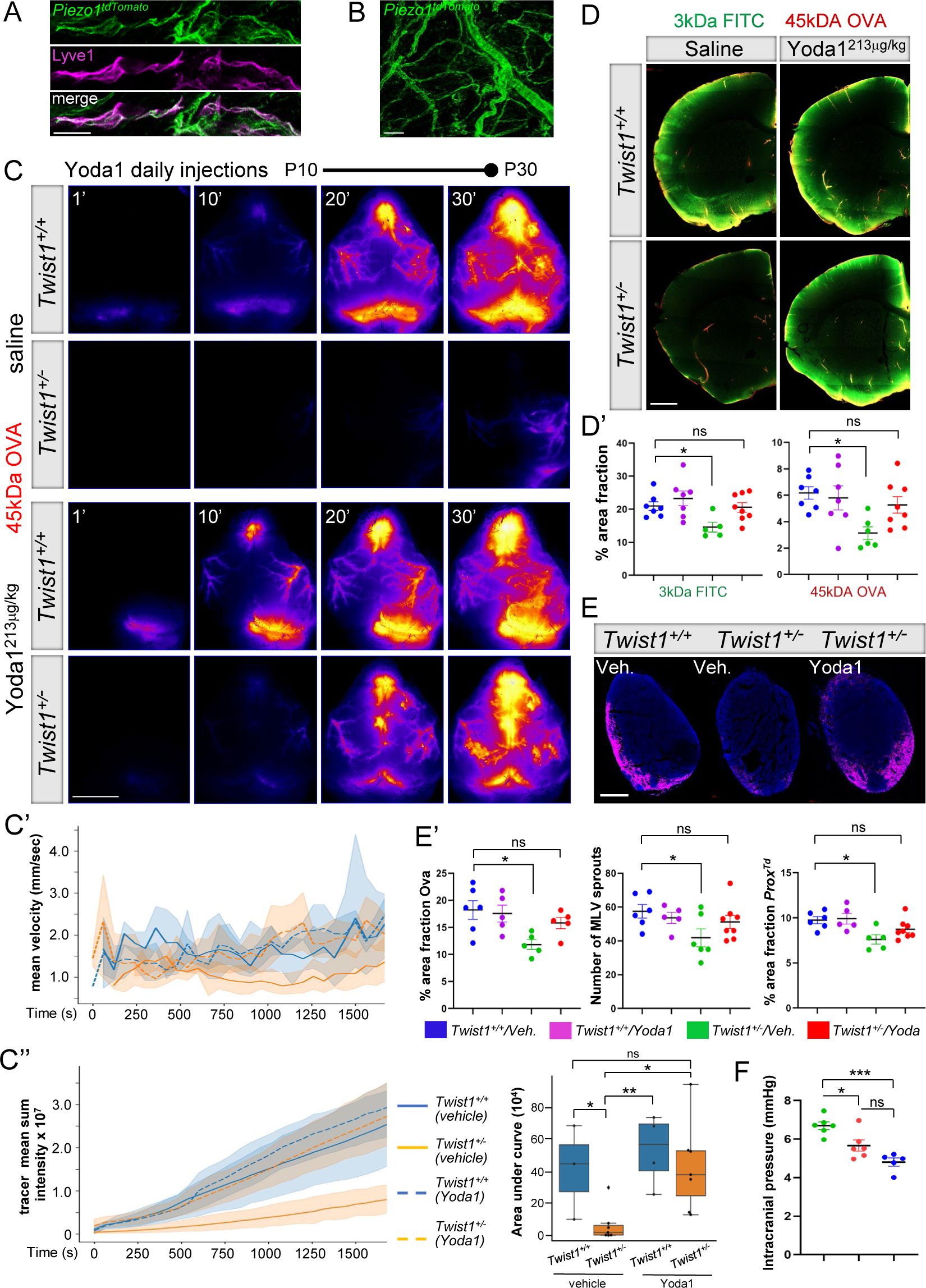
Piezo1 activation helps restore MLV functions and brain-CSF perfusion in CS. (A) *Piezo1^tdTomato^*expression (depicted in green) is detected in MLVs co-labeled with Lyve-1. (B) *Piezo1^tdTomato^* expression is also found in small and large caliber meningeal blood vessels. (C) Representative transcranial timelapse images from juvenile P30 mice treated daily with saline vehicle (top two panels) or 213μg/kg Yoda1 (bottom two panels). Yoda1 helps restore CSF flow along preferred circulation pathways in CS mice. (C’) Graphs depicting tracer mean velocity and (C’’) mean sum intensity in *Twist1^+/+^* and *Twist1^+/−^* mice treated with vehicle or Yoda1 [*Twist1^+/+^*vehicle (n=3); *Twist1^+/+^* Yoda1 (n=4); *Twist1^+/−^* vehicle (n=6); *Twist1^+/−^* Yoda1 (n=7)]. (D) Representative brain sections from juvenile P30 mice treated daily with saline vehicle (left panels) or 213μg/kg Yoda1 (right panels). Perfusion of CSF macromolecules into the brain is significantly improved in *Twist1^+/−^* mice treated with Yoda1. (D’) Quantification of percent area fraction of 3kDa Dextran and 45kDa Ovalbumin in brain slices [*Twist1^+/+^*vehicle (n=7); *Twist1^+/+^* Yoda1 (n=7); *Twist1^+/−^* vehicle (n=5); *Twist1^+/−^* Yoda1 (n=8)]. (E and E’) Representative dCLN sections from juvenile P30 mice treated daily with saline vehicle or with 213μg/kg Yoda1. Yoda1 treatment helps restore ovalbumin drainage to the dCLNs [*Twist1^+/+^*vehicle (n=6); *Twist1^+/+^* Yoda1 (n=5); *Twist1^+/−^* vehicle (n=5); *Twist1^+/−^* Yoda1 (n=5)], the number of MLV sprouts [*Twist1^+/+^* vehicle (n=6); *Twist1^+/+^* Yoda1 (n=5); *Twist1^+/−^* vehicle (n=6); *Twist1^+/−^* Yoda1 (n=8)] and percent area coverage of *Prox1-tdTomato* signal in dura [*Twist1^+/+^* vehicle (n=6); *Twist1^+/+^* Yoda1 (n=5); *Twist1^+/−^* vehicle (n=6); *Twist1^+/−^* Yoda1 (n=8)]. (F) Yoda1 significantly reduces ICP in young adult *Twist1^+/−^* mice with unilateral or bilateral CS [*Twist1^+/−^* vehicle (n=6); *Twist1^+/−^*Yoda1 (n=6); *Twist1^+/+^* vehicle (n=5)]. \**p≤0.05*, *\*\*p≤0.01, ***p≤0.001.* One-way ANOVA with Bejamini-Hochberg correction (C’’), Dunnett’s (D and E), and Tukey’s (F) multiple comparison test. Scale bar: (A) 20μm; (B) 50μm; (C) 5mm; (D) 1mm; (E) 200μm

Yoda1 is a small molecule agonist that has high specificity and affinity for Piezo1, without off-target effects, and lowers the mechanical threshold for channel activation (39). We treated *Twist1^+/−^*mice daily, starting at P10 until P30, with intraperitoneal injections of Yoda1 and then tested if it could restore the perfusion of CSF macromolecules into the brains of CS mice. We injected 3kDa dextran and 45kDA ovalbumin tracers into the cisterna magna and sacrificed the mice 30 minutes later. In juvenile *Twist1^+/−^:Prox1-tdTomato* mice treated with saline vehicles, CSF circulation along dorsal pathways was perturbed compared with unaffected littermates receiving saline vehicles, similar to findings in adult mice (**Figure 7C**). In postmortem tissue, there was also a significant reduction in the amount of both tracers in the brain parenchyma (**Figure 7D, D’**). Strikingly, in *Twist1^+/−^:Prox1-tdTomato* mice treated with Yoda1, CSF flow along dorsal pathways was restored (**Figure 7C-C’’**). Furthermore, perfusion of tracer into the brain was also improved as the differences between *Twist1^+/−^:Prox1-tdTomato* mice treated with Yoda1 and unaffected littermates treated with either saline vehicles or Yoda1 were no longer significant (**Figure 7D, D’**). We still, however, found subtle decreases in AQP4 immunolabeling at glial endfeet abutting larger caliber vessels, which may have possibly prevented a more complete rescue in some animals **(Figure S6D)**.

Next, we tested whether macromolecule drainage to the dCLNs also improved. Owing to the random selection of animals at P10 when the full extent of suture fusion is harder to gauge because it is still an ongoing process (40), MLV phenotypes in P30 *Twist1^+/−^:Prox1-tdTomato* mice treated with saline vehicles were slightly less severe due to milder suture fusion in this particular cohort. Nonetheless, MLV sprouting and coverage along the transverse sinuses plus drainage to the dCLNs was improved in *Twist1^+/−^:Prox1-tdTomato* mice treated with Yoda1, as the differences were no longer significant compared to unaffected controls treated with vehicle or Yoda1 (**Figure 7E, E’**). Thus, activating Piezo1 with small molecule agonists can help rescue insults to meningeal lymphatic functions and brain-CSF perfusion in mice with CS.

We next tested whether improvement to brain-CSF perfusion and meningeal lymphatic functions in *Twist1^+/−^* CS mice could result from reducing the levels of ICP. Treating *Twist1^+/−^* mice that had either unilateral or bilateral fusion with Yoda1 significantly reduced the levels of ICP; ICP was significantly lower compared to CS mice receiving saline vehicle, whereas the differences were no longer significant compared to untreated control mice (**Fig. 7F**). Thus, activating Piezo1 with a small molecule agonist in mice with CS may help rescue insults to CSF flow, meningeal lymphatic drainage, and brain-CSF perfusion, at least in part, by reducing the levels of ICP. These findings are notable as they also suggest that raised ICP in CS may be caused, in large part, from effects exerted by the vasculature and blood flow.

Finally, we asked if Piezo1 activation could also restore MLV growth and sprouting in CS mice. We treated P10 *Twist1^+/−^* mice and littermate controls for seven consecutive days with intraperitoneal injections of Yoda1. In P17 *Twist1^+/−^:Prox1-tdTomato* mice treated with saline vehicle, we again saw less MLV sprouts and growth along the transverse sinus, confluence, and superior sagittal sinus (**Figure. 8A, A’**). However, meningeal lymphangiogenesis was rescued in *Twist1^+/−^:Prox1-tdTomato* mice that received Yoda1 (213ug/kg), as these differences were no longer significant between unaffected littermates treated with saline or those treated with Yoda1 (**Figure. 8A, A’**). Thus, activating Piezo1 signaling in *Twist1^+/−^* CS pups can also rescue impairments to MLV growth and expansion.

**Figure 8:**
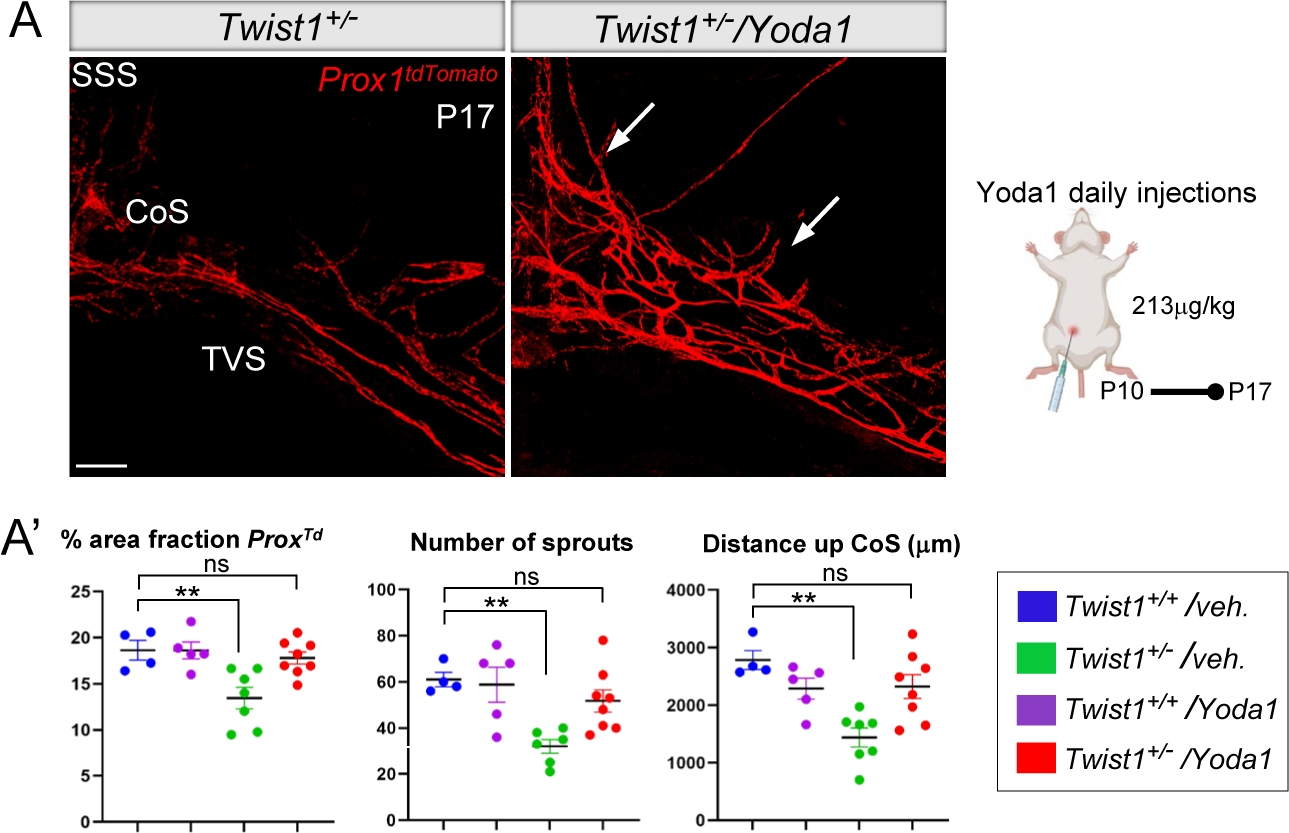
Piezo1 activation restores MLV growth and sprouting in juvenile CS pups. (A) Representative images from juvenile P17 mice treated daily with saline vehicle or 213μg/kg Yoda1. Yoda1 restores MLV growth and sprouting along the transverse sinuses. (A’) Quantification of percent area fraction of *Prox1-tdTomato* signal, number of sprouts, and distance of growth from the confluence of sinuses [*Twist1^+/+^* vehicle (n=4); *Twist1^+/+^* Yoda1 (n=5); *Twist1^+/−^* vehicle (n=7); *Twist1^+/−^* Yoda1 (n=8)]. SSS=superior sagittal sinus; TVS=transverse sinus; CoS=confluence of sinuses. \*\**p≤0.01,* One-way ANOVA with Dunnett’s multiple comparison test. Scale bar: (A) 200μm

### Stimulating Piezo1 in aged animals helps restore MLVs, drainage to the dCLNs, and brain-CSF perfusion

Pathological changes to MLVs, drainage to the dCLNs, and brain-CSF perfusion in CS mice are reminiscent of findings from aged animals, which have diminished MLV drainage and brain-CSF perfusion compared to young and middle-aged adults. We therefore tested whether stimulating Piezo1 mechanotransduction signaling with Yoda1 could also improve MLV drainage and brain-CSF perfusion in mice aged 22-24 months. Yoda1 treated mice showed a significant increase in brain CSF-perfusion as measured by the amount of tracer in brain tissue (**Figure 9A, A’**). Furthermore, Yoda1 treated mice showed a significant increase in MLV complexity, as measured by the number of loops and sprouts along the transverse sinus and especially at the confluence (**Figure 9B, B’**). Vessel coverage along the superior sagittal sinus was also increased, as was the amount of tracer in the dCLNs (**Figure 9B, C, C’**). Thus, stimulating Piezo1 mechanotransduction signaling with small molecule agonists can significantly enhance MLV coverage, drainage, and brain-CSF perfusion in aged mice.

**Figure 9:**
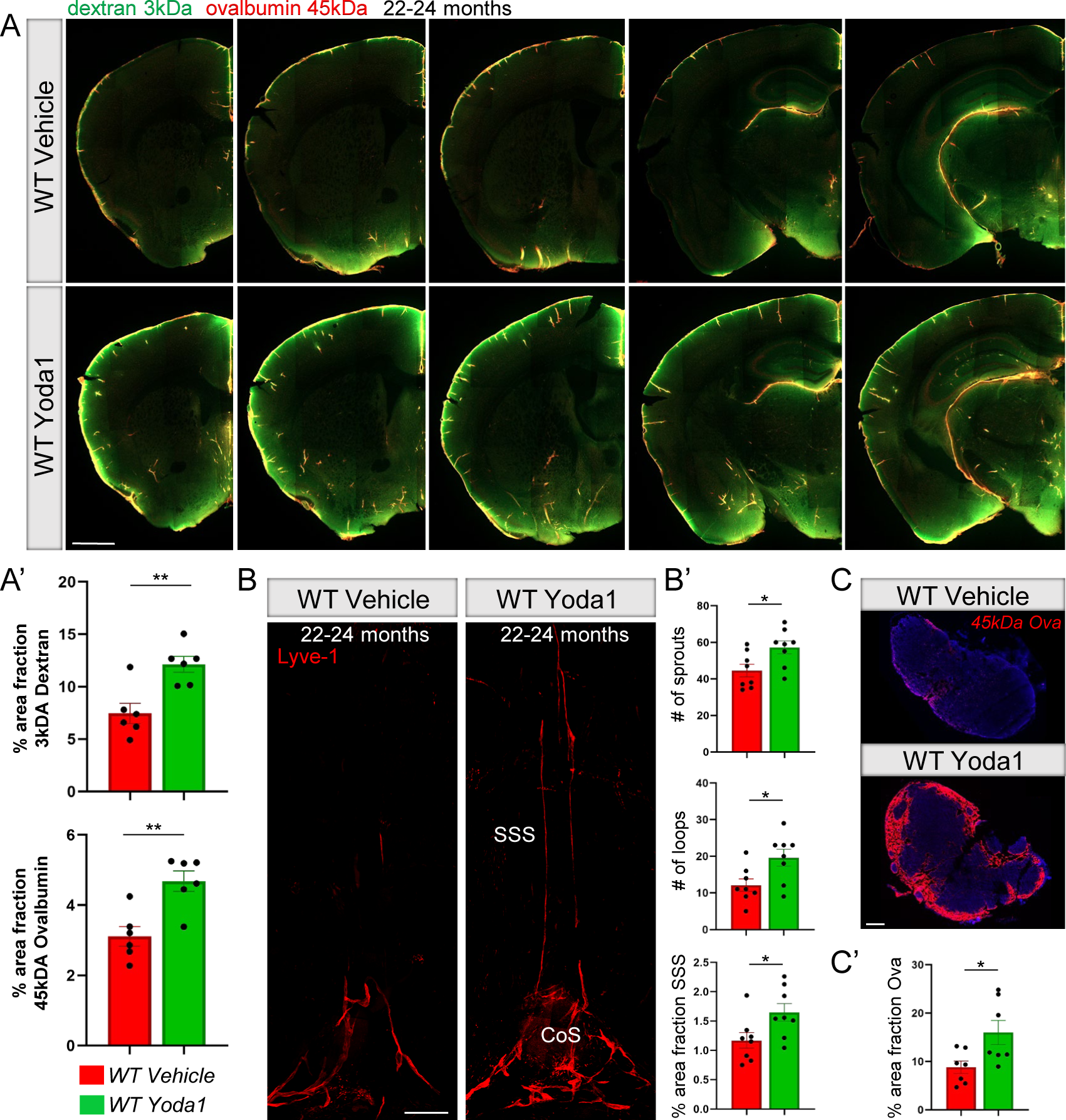
Piezo1 activation helps restore MLV networks and brain-CSF perfusion in aged animals. (A) Perfusion of CSF macromolecules into the brain is reduced in 22–24-month-old mice treated with saline vehicle. Yoda1 treatment improves brain-CSF perfusion as seen by a significant increase in the amount of 3kDa dextran and 45kDa ovalbumin tracer in tissue (A’) [WT vehicle and WT Yoda1 (n=6)]. (B) Yoda1 treatment in 22-24-month-old mice increases MLV coverage along the SSS and the numbers of loops and sprouts along the sinus confluence and proximal transverse sinus. (B’) Quantification of numbers of sprouts, loops and percent area fraction of Lyve-1 signal along the SSS [WT vehicle and WT Yoda1 (n=8)]. (C) Tracer drainage to the dCLNs is significantly improved in 22-24-month-old mice receiving Yoda1 versus saline vehicle. (C’) Quantification of percent area fraction of 45kDA ovalbumin tracer [WT vehicle and WT Yoda1 (n=7)]. SSS=superior sagittal sinus; CoS=confluence of sinuses. \**p≤0.05,* \*\**p≤0.01,* Two-tailed unpaired t test. Scale bar: (A, B) 500μm; (C) 200μm

## Discussion

MLVs were first described in the late eighteenth century by Italian anatomist Paolo Mascagni, but his observations were disputed and difficult to reproduce, obscuring further recognition and investigation of these unique vessels (41). The re-discovery of MLVs more than two centuries later has reignited interest and debate about how these vessels develop within their unique environment, how their physiology and functions compare with traditional lymphatics, and their specialized role for facilitating neuroimmune surveillance and waste clearance from the brain. More genetic models are still needed to permit studies into how MLVs develop, factors that promote their long-term maintenance, and also their support of brain waste clearance. Pathological changes to MLVs in multiple genetic models for CS underscore their utility for elucidating non-cell autonomous interactions with the environment that drive meningeal lymphangiogenesis and the long-term maintenance of lymphatic networks in dura. Furthermore, deficits in brain-CSF perfusion seen in CS models can also provide novel insights into how MLVs function to help support the clearance of macromolecules and neurotoxins from the brain. Finally, our ability to rescue lymphatic growth, drainage, and brain-CSF perfusion in CS with a small molecule agonist for Piezo1–which we showed provides similar benefits in aged mice–provides novel avenues to help restore brain waste clearance and stave off the toxic effects of waste on brain health and function with ageing.

Our findings in multiple genetic models show that CS impairs lymphatic sprouting and expansion in dura. In homozygous *Twist1^FLX/FLX^:Sm22a-Cre* mice, this was previously attributed to loss of growth factor signaling from venous smooth muscle and/or hypoplastic dura (22). However, the venous sinuses and surrounding dura are unaffected in *Twist1^+/−^*, *Twist1^+/FLX^:Sm22a-Cre*, and *Fgfr2^+/S252W^:Sm22a-Cre* mice. Instead, we show that CS causes raised ICP and impedes CSF flow, a consistent finding in all four models, including homozygous *Twist1^FLX/FLX^:Sm22a-Cre* mice. In addition, activating Piezo1, a mechanosensitive ion channel, with Yoda1 significantly reduces the levels of ICP in *Twist1^+/−^*mice, and this is associated with significant improvement to meningeal lymphatic growth, drainage, and also brain-CSF perfusion. Thus, our results now suggest that Piezo1 may be involved in the regulation of ICP. To our knowledge, this is also the first demonstration of a small molecule treatment effective for reducing ICP in CS (or any disorder). Our results also suggest that raised ICP in CS may be, in large part, vascular and fluid based as Yoda1 was able to reduce ICP in mice with malformed skulls and fused sutures. Activating Piezo1 signaling in other fluid disorders that associate with raised ICP may have similar beneficial effects on lymphatic networks and brain-CSF perfusion.

In addition to reducing ICP, it is also possible that Yoda1 has direct effects on MLVs via activation of Piezo1 mechanotransduction signaling. Laminar flow is required for lymphatic growth and maintenance, and it activates Piezo1 mechanotransduction signaling in lymphatic vessels to promote lymphangiogenesis (7, 42, 43). For example, conditional inactivation of *Piezo1* in embryonic lymphatic endothelial cells inhibits their sprouting and expansion, whereas activation of Piezo1 signaling increases lymphatic vessel density and number of vascular tips in three-dimensional cultures (7). In addition, *Piezo1* inactivation in postnatal mice causes lymphatic regression in the mesentery and hindlimb, suggesting laminar flow may be required for the long-term maintenance of MLVs. Thus, Piezo1-mediated mechanotransduction signaling supports lymphatic sprouting, network expansion, and the maintenance of pre-existing networks, processes that all appear to be affected in CS mice. Another intriguing possibility is that CSF flow provides mechanical force to facilitate MLV sprouting and expansion, and limiting MLV access to CSF impairs their development and long-term maintenance in CS, potentially via perturbing Piezo1 signaling and lymphatic mechanotransduction signaling pathways. Notably, MLVs lining meningeal arteries are not affected in CS mice and these vessels do not appear to have access to CSF, in agreement with previous studies (5). Thus, the improvement to CSF circulation following reduction of ICP through Yoda1 treatment may, in turn, exert mechanical force that facilitates the restoration of meningeal lymphatic growth and drainage. However, it is important to emphasize that further studies, including the effects of lymphatic-specific knockout of Piezo1 on MLVs, are still needed, as this was not directly addressed in the current study. Furthermore, increasing interstitial flow and functional drainage can stimulate VEGF-C expression, VEGFR3 activation, and also cell proliferation via integrin-dependent interactions between lymphatic endothelial cells and the extracellular matrix (8). Thus, it remains possible that multiple signaling pathways, including BMP signaling, may be implicated in CS mice, affecting lymphatic vessels. This will require further investigation.

Our data in aged mice also suggest that CSF may support long-term maintenance of MLVs, and the natural decline of CSF flow and turnover as animals age may cause these vessels to regress or undergo pathological changes that are indicative of functional impairments to fluid drainage. Similar to peripheral lymphatics, MLVs require VEGF-C/VEGFR3 signaling for their initial growth and expansion (2). Once networks have fully formed, however, VEGF-C/VEGFR3 signaling is still required for their survival and long-term maintenance, particularly for dorsal vessels along the transverse and superior sagittal sinuses that are very sensitive to the levels of VEGF-C (2). Similarly, our data in aged mice now suggest that mechanotransduction signaling in MLVs via Piezo1 may promote sprouting and expansion and also long-term maintenance in adults, presumably more likely due to direct effects on vessels versus changes to ICP (which is not known to be affected in aged but otherwise healthy mice). The gradual decline of CSF flow, turnover, and drainage to the dCLNs as humans and mice age, or changes to preferred circulation pathways in pathological states, may inhibit both VEGF-C/VEGFR3 signaling and Piezo1 activation, causing dorsal networks to regress and basal networks to undergo lymphedematous-like changes. Considering adenoviral-mediated expression of VEGF-C can improve MLV functions and help restore brain-CSF perfusion in aged animals–similar to the effects of Yoda1–combined treatments may provide greater beneficial effects for the long-term functional maintenance of MLVs and brain-CSF perfusion (17).

The demise of brain waste clearance systems with ageing or in pathological states is proposed to be an underlying risk factor for dementia (44, 45). Brain-CSF perfusion is most active during sleep (46, 47), and sleep disorders are common in the elderly. As mice age, dorsal MLVs begin to regress whereas basal MLVs become hyperplastic, concurrent with the decline of brain-CSF perfusion (16, 17, 44). As such, the demise of brain waste clearance systems in CS appears to have early onset. Initially, the growth and sprouting of MLVs is affected in juveniles. In young adults, MLV networks are typically hypoplastic but some animals with milder CS show less affection, and a small subset may instead show signs of vessel hyperplasia. As mice reach middle age at 8-12 months, dorsal MLVs have continued to regress and basal vessels become hyperplastic. The distinct pathological changes to dorsal versus basal MLVs may reflect their exposure to different flow conditions in CS, which also change with ageing as demonstrated by contrast-enhanced magnetic resonance imaging (14). Furthermore, the natural decline of CSF flow and turnover with ageing may be comparable to the loss of normal CSF circulation as seen in CS mice. These data suggest impairments to CSF flow in pathological conditions such as CS may accelerate the deterioration of MLVs and brain waste clearance. Thus, genetic models for CS have the power to elucidate not only how these systems develop, but also the factors and signaling mechanisms that are required for long-term maintenance.

Brain-CSF perfusion and clearance of interstitial solutes and neurotoxins from the brain requires AQP4 water channels at glial endfeet, which line paravascular spaces surrounding blood vessels (35). In pharmacological models of MLV ablation using Visudyne treatment, brain-CSF perfusion is significantly reduced without changes to AQP4 localization or expression (17). In juvenile CS mice at P17, we see an initial delay in AQP4 expression and polarization to endfeet. However, we do not see wide-spread changes to AQP4 expression or localization in adult CS mice with significantly impaired brain-CSF perfusion, with the exception of subtle reductions at glial endfeet along larger caliber vessels. Considering AQP4 knockout mice show only a 30-50% reduction in the perfusion of CSF macromolecules into the brain, the changes we detect in CS mice (∼10-20% reduction of AQP4 along larger caliber vessels) are unlikely to significantly affect brain-CSF perfusion, although we cannot rule out some minor effects. Interestingly, as mice age, AQP4 is lost at glial endfeet that line penetrating pial arterioles (44), in line with our observations that brain waste clearance systems show signs of precocious ageing in CS. Overall, our data agrees with previous reports that suggest brain-CSF perfusion is integrated with lymphatic drainage in the dura (17), and ageing-related or pathological conditions that restrict drainage to the dCLNs are likely to have deleterious consequences for brain-CSF perfusion and waste clearance. It is also important to note that brain-CSF perfusion is dependent upon cerebral arterial pulsatility, which declines in aged mice that have glymphatic deficits (44). Improved brain-CSF perfusion in aged mice following Yoda1 treatment may also involve changes to arterial pulsatility and blood flow, an avenue for future investigation in both aged mice and those with CS.

Combined deficits in MLV drainage and brain-CSF perfusion are predicted to exacerbate amyloid-beta pathology in *Twist1^+/−^:5xFAD* models with CS. Our findings agree with previous studies that show restricting MLV drainage and brain-CSF perfusion increases amyloid-beta burden in the hippocampus of *5xFAD* mice (17). Furthermore, ablating MLVs leads to worse outcomes and greater deposition of amyloid-beta in *5xFAD* mice treated with anti-Aβ passive immunotherapy, whereas increasing MLV functions with exogenous VEGF-C improves efficacy (18). Interestingly, recent findings in *5xFAD* mice show that intraperitoneal injections of Yoda1, which is able to cross the blood-brain barrier, can increase amyloid-beta phagocytosis via effects on microglia that express Piezo1 (48, 49). It remains to be seen what effects Yoda1 has on amyloid-beta burden in *Twist1^+/−^* :*5xFAD* mice. However, our results suggest Yoda1 may act to reduce amyloid-beta burden in *Twist1^+/−^:5xFAD* mice through combined effects on microglia and mechanosensing, as well as improving MLV drainage and brain-CSF perfusion. Moreover, Yoda1 may provide similar benefits to counteract the buildup of neurotoxins, such as tau and amyloid-beta, with ageing and in response to brain injury.

Deficits in MLV drainage and brain-CSF perfusion in CS show how skull expansion is integrated with the development and maintenance of brain waste clearance systems. The human skull expands rapidly after birth, reaching ∼80% of its adult size by year two (approximately ∼2-3 weeks in mice) (50). Concurrently, the brain nearly triples in size and, along with increasing volumes of blood and CSF within the fixed confines of the skull, this causes a sharp rise in the levels of ICP (51). ICP needs to be controlled in order to maintain proper fluid dynamics in the head. This suggests skull expansion, via effects on ICP and CSF, exerts non-cell autonomous control over MLV development. Also, the growth and remodeling of MLVs along the venous sinuses mirrors the basal-dorsal establishment of CSF circulation pathways while the skull rapidly expands in mice (4). As venous sinuses and MLVs have access to CSF (9, 10), impediments to CSF flow caused by CS can potentially affect MLVs by restricting their access to CSF. Interestingly, arterial MLVs are not affected in CS and we do not detect tracer around these vessels, suggesting their development and functions are less dependent on CSF and perhaps more related to draining dural ISF. In addition, the polarization of AQP4 water channels to glial endfeet also mirrors the establishment of CSF circulation pathways (4). It is therefore possible that both the development and maintenance of the glymphatic and meningeal lymphatic systems may be interconnected via shared dependency on CSF flow and mechanical forces that are influenced by skull growth. Thus, failure to maintain normal CSF flow in conditions like CS or hydrocephalus is likely to affect MLV development and brain-CSF perfusion, potentially leading to the accumulation of waste and neurotoxins.

Cognitive impairments and learning disabilities are recognized in CS (52). The underlying causes are still unclear in most cases, although raised ICP is associated with neurological and cognitive dysfunction in CS (24). Surgical interventions to alleviate raised ICP include removing fused sutures and invasive craniotomies to remodel and expand the skull. While these surgeries help improve airways and cosmetic features, they sometimes fail to maintain normal levels of ICP. For example, upwards of 40% of individuals with *TWIST1* mutations and CS may still experience lingering, raised ICP following surgery (53). Interestingly, a recent study showed that engraftment of sutural progenitor cells into the affected sutures of *Twist1^+/−^* mice with bilateral synostosis prevented fusion, improved skull growth, and alleviated raised ICP and cognitive impairment (26). Cognitive deficits were attributed to cortical thinning, but these findings were not examined in mice with unilateral fusion, which is common and also impedes CSF flow in our models. Our results now suggest deficits in MLV drainage, CNS waste clearance, or even altered neuroimmune surveillance may contribute to cognitive impairment in CS. Furthermore, those who suffer from CS and/or chronically raised ICP may have increased risk for neurodegenerative disease. Our novel findings showing that Yoda1 can reduce ICP and help restore MLV functions and brain-CSF perfusion in adult CS mice therefore has important clinical implications, as similar treatments may hold promise in humans. Furthermore, it will also be important to test whether surgical and/or regenerative medicine approaches that alleviate ICP in CS can repair MLVs and restore normal brain-CSF perfusion.

## Materials and Methods

### Mice

Mice were maintained on a normal light/dark cycle and experiments were performed during the light cycle. The following transgenic mice were used: *Twist1^FLX^* (RRID:MMRRC_016842-UNC), *Prox1^tdTomato^* (RRID:MMRRC_036531-UCD), *Sm22a-cre* (RRID:IMSR_JAX:017491), *Sox2-cre* (RRID:IMSR_JAX:008454), *Fgfr2^S252W^*(kind gift from Ethylin Jabs, Mt. Sinai), *5xFAD* (RRID:MMRRC_034840-JAX), and *Piezo1^tm1.1Apat/j^* (RRID:IMSR_JAX:029214). Male and female mice were included for all experiments, with the exception of the aged cohorts that received saline or Yoda1 treatments, of which only female animals were used (RRID:IMSR_JAX:000664). Animals were maintained on C57Bl/6 or a mixed genetic background (C57Bl/6:CD1). *Twist1*^+/−^ mice were generated by mating *Twist1^FLX/+^* males to female *Sox2*-cre mice in order to achieve germline recombination. The ages of animals in this study include postnatal days (P) 17, 30, 2-4 months of age, 6-8 months of age, 10-14 months of age, and 22-24 months of age. For postnatal day 17 (P17) timepoints, mutant mice were only included if they exhibited obvious synostosis. Adult animals with CS were included if they exhibited full unilateral or near-complete/full bilateral coronal synostosis, as judged by visual inspection under a stereomicroscope. Animals with unilateral fusion were also included if greater than 50% of the suture was fused and the skull was dysmorphic, as shown in **Figure 1B**. Examples were chosen for further analysis using X-ray microscopy. Aged animals were purchased from JAX labs at 96 weeks and were allowed to mature to 22-24 months in our colony.

### Antibodies

The following antibodies were used: rabbit anti-Lyve1 (1:500, Abcam ab14917 RRID:AB_301509), rabbit anti-RFP (1:1500, Rockland, 600-401-379, RRID:AB_2209751), rabbit anti-amyloid beta (1:400, Cell Signaling, D54D2, RRID:AB_2797642), rabbit anti-Aqp4 (1:500, Invitrogen PA5-78812, RRID:AB_2745928), rabbit Mouse anti-Crabp2 (1:100, Millipore, MAB5488, RRID:AB_2085470), mouse anti-smooth muscle actin (1:500 Sigma, C6198, RRID:AB_476856), GS-Lectin IB4 488 and 649 (1:100, Vector Biolabs, #B-1205-.5 and #DL-1208-.5), chicken anti-IBA1 (1:500, Aves, IBA1-0100, RRID:AB_2910556), mouse anti-GFAP (1:500, Santa Cruz, SC33673

### Immunohistochemistry

Following skull decalcification, tissue was stained overnight in primary antibody solution containing 5% NGS in PBS with 0.3% Triton. Secondary antibodies were diluted 1:1000 in PBS with 5% NGS and 0.3% Triton and applied at room temperature for 1 hour. For AQP4 staining, fixed 100μm brain sections were incubated in Dent’s Fix (80% MeOH, 20% DMSO) overnight at 4°C prior to primary antibody staining. Piezo1 labeling in Piezo1^tdTomato^ mice was performed as previously reported (54). Briefly, mice were perfused with PBS containing heparin, and the dura was immediately dissected from the skull and fixed in 4% PFA for 20 minutes. The tissue was quenched using 20mM glycine and 75mM ammonium chloride with 1% triton X-100 in PBS for 20 minutes. The tissue was washed in PBS and then incubated in blocking buffer (0.6% bovine skin gelatin with 0.05% saponin in PBS with 5% normal donkey serum) for one hour at room temperature. Tissue was stained with anti-rabbit RFP antibody and anti-rat Lyve-1 overnight at 4°C in blocking buffer without serum. The tissue was washed, stained with appropriate secondary antibodies, and flat mounted onto a slide for imaging.

### Dorsal and Basal Skull Flat mounts

For dorsal skull flat mounts, dorsal craniotomies were performed by making an incision at the foramen magnum and using curved tip spring scissors to cut radially around the base to the orbits. Then a final incision was made cutting the bone between the two orbits, freeing the skull cap from the rest of the cranium. Skull caps with meninges attached were postfixed in 4% PFA overnight with agitation at 4°C. After fixation, skull caps were incubated in 3% Hydrogen Peroxide for 24 hours at 4°C, followed by Dent’s Fix overnight at 4°C (80% MeOH, 20% DMSO). Skulls were decalcified in 14% EDTA for 3-5 days with solution changes every other day until the skull was malleable. Dorsal skull preparations were then cut along the transverse and superior sagittal sinuses, the remaining excess bone and the temporal bone were removed, and the tissue was covered with 600 µL of mounting media before being cover slipped.

### Skull 3D X-Ray Microscopy (computed tomography)

Animals were sacrificed via transcardial perfusion with 4% PFA. The heads were then decapitated and post-fixed overnight in 4% PFA. Hair and skin were removed prior to imaging. Images were obtained using a Bruker Skyscan 1272 X-ray microscope. The following scan conditions were used: image pixel size=13.5 μm, camera=1632 columns×1092 rows, rotation step=0.4 degrees, frame averaging=3, filter=1 mm Al. The resulting images were reconstructed and converted to dicom format with Skyscan Ctan software. Dicom files were opened in Vivoquant for segmentation of teeth and bone from less dense soft tissue.

### Intracranial pressure (ICP) recordings

Animals aged between two to four months were placed on stereotaxic frame and anesthetized with a constant supply of 1% Isoflurane and 2% oxygen. Protocol was adapted from Da Mesquita et. al, 2018. A midline incision was made from just caudal to the orbits to the apex of skull vault. The underlying skull was then cleaned with sterile saline. Using a dental drill, a 0.5mm hole was drilled into the right parietal bone (ML: +1.0mm, AP: −1.0mm from bregma, or approximate coordinates in animals with suture fusion). A pressure sensor (Millar SPR100) was inserted perpendicularly at a depth of 1mm. The pressure sensor was connected to a PCU-2000 pressure control unit (Millar). Following placement of the pressure sensor, the waveform was allowed to stabilize and readings were averaged over 5 minutes from the time of signal stabilization to achieve a pressure measurement. For determining the effect of Yoda1 on ICP, adult *Twist1*^+/−^ and wildtype counterparts received intraperitoneal Yoda1 injections (1mg/kg) five times per week for 4 weeks. Adult injections started at one month of age, and at the end of the 4 weeks ICP was measured as previously described. Following recordings, all animals were euthanized.

### Cisterna Magna tracer infusions

Animals were anesthetized using ketamine/xylazine (100 mg/kg). An incision was made on the midline of the head at the occipital crest. Once the skin was excised, curved forceps were used to break through the superficial connective tissue to reveal the underlying muscle. The muscle was carefully separated along the midline to expose the opening to the cisterna magna, and care was taken to not induce tears or bleeding. A 45kDa ovalbumin-647 tracer (Molecular Probes, Invitrogen) and 3kDa fluoresceine dextran tracer (Molecular Probes, Invitrogen) were mixed in artificial cerebrospinal fluid at 2µg/ml and 0.5µg/ml respectively. 5µl of solution was loaded into a 10µl Hamilton syringe attached to polyethylene tubing. The solution was injected using a 28-gauge needle and a Nanomite infusion system (Harvard Apparatus) with an injection rate of 2.5 µl/min. Animals were kept on supplemental oxygen throughout the duration of the experiment to stabilize breathing and minimize hypercapnia, and the needle was glued in place to prevent depressurization. A second dose of ketamine/xylazine (100 mg/kg) was applied approximately 30 minutes after the first dose to ensure animals were properly anesthetized, monitored by tail pinches. After completion of the experiment, the needle was removed and the animals were immediately euthanized by transcardial perfusion. All tracer infusion experiments were 60-minute injections with the exception of P30 Yoda1 experiments, which were performed for 30 minutes.

### Cervical lymph node imaging and quantification

Animals were perfused with 4% PFA and the deep cervical lymph nodes were dissected under fluorescence. The tissue was allowed to post-fix for 12 hours in 2% PFA, prior to sinking in 30% sucrose and embedding in Neg-50 medium. 20µm sections were cut from the entire length of the lymph node. The sections were imaged using a Leica M165FC stereomicroscope equipped with a 1X Plan objective and DFC7000T camera. Ten 20 µm sections per animal from the deep cervical lymph nodes were imaged, equally thresholded, and analyzed by tracing out the tissue sections and calculating the percent area fraction of the ovalbumin 45kDa tracer. Values were averaged for each animal to obtain a single value representing the average percent area fraction of tracer. Representative images were imaged using a LSM800 confocal microscope equipped with a 20x 0.80 NA objective.

### Transcranial live imaging

Mice were prepped for tracer infusion into the cisterna magna, as described above. The head was shaved and dermis peeled back to expose the skull. Mice were kept on supplemental oxygen throughout the duration of the experiment. Mice were positioned and immobilized by placing the nose into the nosecone piece mounted on a metal stage. Dorsal images of the skull (top-down) were captured every minute using a Leica M165FC stereomicroscope equipped with a 1x Plan objective, LED fluorescence, a Cy5 filter, and a DFC7000T camera. Representative images at select time points were converted to 8 bit, uniformly adjusted for intensity, and pseudo-colored using look-up tables (Fire) in Image J. Velocity measurements for CSF tracer at the migrating front were calculated using FrontTracker.m in MATLAB (55). This software calculates the local velocity of several points at the leading edge of the migrating tracer front (by determining the distance the front advances between images) after setting user-defined intensity thresholds, as also described in (4). The mean sum intensity was calculated using the total mean sum fluorescence of tracer at each time point between controls and affected littermates. Following imaging, the animals were immediately euthanized by transcardial perfusion in order to obtain brain tissue and skull caps.

### Imaging and quantification of brain CSF-perfusion

Following transcardial perfusion with 4% PFA, brains were dissected and post-fixed for 24 hours in 4% PFA. They were then embedded in 3% agarose and sectioned (100µm) on a Leica VT-1000 vibratome. To minimize tracer fading, tissue was immediately mounted and imaged on an LSM800 confocal microscope with a 5x 0.16 NA objective. Z-stacks were obtained from a total of six sections ranging from bregma coordinates +1.2 to −2.5. The percent area fraction of tracer was calculated from maximum intensity z-stacks (10μm step size). Whole-brain sections were uniformly thresholded and analyzed using ImageJ. Threshold values for 45kDa OVA-647 were 10 and values for 3kDa dextran were set at 28. Values across brain sections were averaged to produce a single data point for each animal.

### Meningeal lymphatic imaging and quantification

Following the preparation of dorsal or basal skull mounts, tissue was imaged using either a LSM700 confocal equipped with a 10x 0.45NA objective or LSM800 confocal equipped with a 10x 0.45 NA objective. Z-stacks were acquired using 10µm interval step sizes. Following imaging, raw files were z-projected and loaded into ImageJ. Sprouts were counted manually and considered as any blunt-ended LYVE1 or Prox1-positive vessel branching off of the primary lymphatic tree. Vessel diameters along the TVS and hotspot regions were obtained by measuring the width of the vessels along ten different defined regions and averaging those together to receive one representative data point per animal. For P17 timepoints, the growth distance from the confluence of sinuses was quantified by measuring the distance from the bottom of the confluence to the highest region where MLVs were present. For both percent area coverage of vessels as well as OVA-647 tracer, a single ROI was generated encompassing both transverse sinuses and also the confluence of sinuses. The ROI was placed over the vessels in ImageJ, images were uniformly thresholded, and the percent area fraction was measured. The same ROI was used for all adult timepoints, and a separate ROI was used for P17 timepoints.

### Amyloid-β coverage in 5xFAD mice

Mice were fixed by transcardial perfusion with 4% PFA. Brains were dissected from the skull and post-fixed in 4% PFA O/N at 4°C. Brains were embedded in 3% agarose and cut into 100μm thick sections using a VT1000S vibratome (Leica). Sections were incubated O/N in primary antibody solution containing 5% NGS and 0.3% triton, followed by one additional O/N incubation in secondary antibody solution. Maximum intensity z-stacks were obtained using 10μm step sizes and a LSM800 confocal microscope (Zeiss) equipped with a 10x 0.3 NA objective. Images were uniformly thresholded and the percent area fraction of amyloid-β was calculated separately in the hippocampus and dorsal cortex, with a boundary set at the insular cortex. Plaque number was calculated using the ImageJ ‘analyze particles tool, and counting puncta at least 20μm in size in order to minimize background noise.

### AQP4 Immunofluorescence imaging and quantification

100μm sections were incubated in Dent’s Fix (80% methanol, 20% DMSO) and placed on a rocker for 60 minutes at room temperature to improve antibody penetration. Sections were then co-stained with 1:500 rabbit anti-AQP4 primary antibody and 1:100 GS-lectin 488 conjugate in 5% normal donkey serum (NDS) and 0.3% PBS-triton with 1mM calcium chloride (PBS-T + 1 mM CaCl_2_). Sections were incubated overnight at 4°C. Following PBS washes, sections were stained with 1:500 donkey anti-rabbit 546 (cat# A10040, ThermoFisher Scientific) secondary antibody in 5% NDS and 0.3% PBS-T + 1 mM CaCl_2_ for one hour at room temperature. Following secondary antibody staining, the tissue was washed with PBS and stored in primary antibody solution (without AQP4) at 4°C until imaging to preserve the lectin signal. Sections were imaged with a LSM800 microscope and a 10x 0.45NA objective. Z-stacks were obtained with a 10μm step size. Vessels were randomly selected from the dorsal cortex and uniformly thresholded, and each section typically had 8-14 large caliber vessels suitable for analysis. ROIs were selected for single vessels, ensuring at least 95% coverage for lectin. Measurements were obtained by dividing percent area coverage for lectin by the percent area coverage for AQP4. For P17 mice, 10 vessels were selected from a section at the level of the hippocampus. Otherwise, 20 vessels were imaged from two sections, one at the level of the striatum and another at the hippocampus (10/section).

### IBA1 and GFAP staining

Adult *Twist1^+/−^* and wildtype counterparts (5-6 months of age) were fixed by transcardial perfusion with 4% PFA. Brains were dissected from the skull and postfixed overnight in 4% PFA overnight (O/N) at 4°C. Brains were embedded in 3% agarose and sectioned coronally at 100μm. Sections were stained overnight at 4°C with chicken anti-IBA1 (1:500) and mouse anti-GFAP (1:500) in 5% normal goat serum and 0.3% triton in PBS. Following staining, sections were imaged with an LSM800 confocal microscope with z-steps of 10μm and a 20x objective (0.8 NA). Images were z-projected and the Iba1 positive cells were counted using FIJI software. Percent area fraction of Iba1 and GFAP1 on whole brain sections was also calculated using Fiji software. For morphometric analysis, compressed z-stacks were acquired using a 20x objective (0.8 NA) at 1μm step sizes. Images were z-projected and analyzed using FIJI software. Images were first despeckled and then the unsharp mask filter was applied. The image was despeckled again and then equally thresholded (min:40, max:255). Three random regions of interest (ROIs) were drawn and cropped. The number of cell bodies were manually counted in each ROI. The diameter of each soma in every ROI was measured by using the line tool and measuring the soma’s longest length. The "analyze particles" function (size: 5-infinity) was then used on each ROI with a mask shown. The masks were then skeletonized and analyzed using "analyze skeleton 2d-3d." Average endpoints per soma, average junctions per soma, and average branch length per soma were calculated by adding the respective values and dividing by the number of soma in each region of interest for each ROI, which were averaged to obtain one value/mouse.

### Yoda1 Injections

Yoda1 (Sigma SML1558) was dissolved in DMSO (0.568 µg/µL) and then diluted to the final working concentration in sterile saline (3:80 dilution). DMSO dissolved 3:80 in saline was considered vehicle for all Yoda1 control experiments. Mice were weighed daily and Yoda1 or vehicle was injected intraperitoneally at 213µg/kg at 6PM each day for the duration of experimentation (7 days for P17 mice, 20 days for P30 mice, 6 weeks for 22–24-month-old mice). Prior to perfusion fixation or in vivo tracer injections, animals received an additional dose of Yoda1 or vehicle 1hr prior to experimentation. Both Yoda1 and saline vehicle treated animals were healthy with no changes to body weight. Animals were excluded if suture fusion was not present upon visual inspection (n=2).

### Cell fractionation and western blotting

Blood vessel fractionation from whole brain tissue was performed using previously published protocols (56). Briefly, brains were minced with a scalpel in HBSS supplemented with 10mM HEPES (Buffer 1). Samples were homogenized using a long 21 G cannula (0.8 mm × 120 mm) mounted on a 2 mL syringe by aspirating and pushing out several times. The homogenates were centrifuged at 2000 x *g* for 10 min at 4°C. The supernatant, containing the parenchyma fraction, was gently pipetted off and mixed with an equal volume of 2x RIPA buffer before freezing on dry ice and storing at −80°C. The pellet containing blood vessels and tethered glial endfeet was resuspended by sonication (5 sec on/off, 30% amplitude) in 2 mL of buffer 1 with 18% (*w*/*v*) dextran, M_r_ ∼ 70,000) and centrifuged at 4400× *g* and 4 °C for 15 min. Brain vessels were collected on a 20µm cell strainer (pluriSelect, Leipzig, Germany) by centrifugation at 200× *g* for 1 min in a swinging bucket rotor. Vessels were washed twice by resuspending in 1 mL buffer 1 plus 1% (w/v) BSA (buffer 2) on the strainer followed by centrifugation. The purified vessels were rinsed in buffer 2 on the strainer, transferred to a 1.5 mL tube, and sedimented at 2000× *g* at 4°C for 5 min. To remove BSA, the resulting pellet was resuspended in 1 mL buffer 1, followed by centrifugation at 2000× *g* at 4°C for 5 min. The supernatant was discarded and the pellet containing blood vessels was frozen on dry ice and stored at −80°C. The vascular fraction (2μg/sample) was run on 4-12% Bis-Tris gels (NuPAGE) according to manufacturer’s instructions and transferred to PVDF membranes. Membranes were blocked with 1X TBS, 0.1% Tween-20 with 5% w/v nonfat dry milk and blotted with anti-rabbit AQP4 (Millipore #AB3594). Following incubation with primary antibody, the blots were stripped (Abcam ab282569) and incubated with mouse β-actin (Sigma #A2228, 1:2000). Following washes, secondary sheep anti-mouse IgG-HRP (Sigma #AC111P) was added in blocking buffer. Gels were incubated in chemiluminescence agent and blots were exposed using film. Protein was quantified from three different biological replicates (whole brains) representing unaffected controls and also affected littermates. Quantification was performed using Image J by placing rectangular boxes on each band and using the Analyze Gels function. The areas under the peaks for AQP4 and the β-actin loading control were marked off and selected using the wand tool. The final relative quantity for each sample was determined by calculating the ratio of the sample band by the loading control band.

### Statistics

Quantifications were performed with investigators masked to genotypes. All statistical analyses were performed using GraphPad Prism software. In general, all studies with two comparison groups were analyzed with unpaired 2-tailed t test, unless otherwise noted. Experiments with more than two groups were analyzed with one way ANOVA with Dunnett’s or Tukey’s multiple comparisons test, unless otherwise noted. Show the median (horizontal line), the 25^th^ and 75^th^ percentiles (boxed region), and min/max data points. All data are expressed as mean ± SEM. All p values < 0.05 were considered significant.

### Study approval

Mouse strains, breeding, and experiments were approved by Rutgers IACUC under protocol PROTO201702623 (MAT).

### Data Availability

Data is available from corresponding author upon request

## Author contributions

Conceptualization: M.J.M., P.S.A., G. B., M.A.T.; Formal analysis: M.J.M., A.J., A.R., J.K.T., M.A.T.; Data curation: M.J.M., with some assistance by P.S.A., J.W., A.J., A.K.; Technical assistance and reagents: Y.H., C.C.S.; Writing - original draft: M.A.T.; Writing - review & editing: M.J.M., A.K., A.J., M.A.T; Supervision: M.A.T.; Funding acquisition: M.A.T.

## Acknowledgements

The authors thank Ethylin Jabs, Mt. Sinai, for providing *Fgfr2^S252W^* mice; the Rutgers Molecular Imaging Center (D. Adler and P. Buckendahl) for assistance with skull 3D x-ray microscopy using the Skyscan 1272 micro-CT scanner (funding by NSF Major Research Instrumentation Award 1828332); Garvit Gupta for assistance with mouse transcranial imaging. Medha Pathak for providing *Piezo1^tdTomato^* mice. A license for Biorender (M.A.T) was used to create schematics in **figure 4C**, using illustration ‘*mouse skull, with mandible and brain*.’

