## Supplemental Figures for "Piezo1 agonist restores meningeal lymphatic vessels, drainage, and brain-CSF perfusion in craniosynostosis and aged mice"

**Supporting Material**

**Figure S1: Dura mater and the venous sinuses are unaffected in *Twist1*$^{+/FLX}$*:Sm22a-Cre* and *Twist1*$^{+/-}$ CS models, and MLVs along the PPA and MMA do not have direct access to CS and develop normally.** (A) Dural scraping from a six-month-old *Twist1*$^{+/-}$ CS mouse (n=3) with regressed dorsal MLVs. The transverse sinuses are both present. Smooth muscle coverage is normal on the transverse (A') and superior sagittal sinuses (A"). (B) Coronal cross section at the dorsal midline of a P0.5 mouse labeled with Crabp2, which marks dura and the underlying arachnoid. The dura and arachnoid membranes show normal Crabp2 expression and are not hypoplastic in *Twist1*$^{+/-}$ CS mice. (C) Dorsal craniotomy and dural flat mount. The dura is easily peeled off the skull in *Twist*$^{+/-}$ and *Twist1*$^{+/FLX}$*:Sm22a-Cre* mice (shown). Representative images from young adult mice aged 2-4 months. (D and D') The percent area fraction of *Prox1-tdTomato* signal along the MMA is normal in *Twist1*$^{+/-}$ CS mice compared with unaffected littermates [*Twist1*$^{+/+}$ (n=3); *Twist1*$^{+/-}$ (n=3)]. Note that tracer is present (arrowhead) in perisinusoidal dura enveloping the PSS (petrosquamosal sinus) but is absent in dura surrounding the PPA (pterygopalatine artery) and MMA (middle meningeal artery). (E) Schematic overview of dorsal and basal perisinusoidal MLVs in mouse. ns, non-significant, *two-tailed unpaired t test.* Scale bar: (A) 250$\mu$m; (B) 50$\mu$m; (D) 1mm.

**Figure S2: MLV networks, flow to perisinusoidal dura, drainage to the dCLNs, and brain-CSF perfusion are unaffected in *Twist1*$^{+/-}$ *mice* without suture fusion.** (A) Representative images show that dorsal MLV networks, stained with Lyve-1, and hotspots along the transverse sinus (TVS) develop normally in *Twist1*$^{+/-}$ mice that lack coronal suture fusion. (A') Following injection of 45kDa ovalbumin tracer into the cisterna magna, tracer deposition in perisinusoidal dura surrounding MLVs and hotspots is comparable between controls and *Twist1*$^{+/-}$ mice that lack fusion of the coronal sutures, versus *Twist1*$^{+/-}$ mice that have coronal suture synostosis. (B) Quantification of percent area fraction Lyve-1 and ovalbumin 45kDa [*Twist1*$^{+/+}$ (n=4); *Twist1*$^{+/-}$ no fusion (n=5); *Twist1*$^{+/-}$ fusion (n=6)]. (C) Skull overviews showing patent coronal sutures (CorS) in a *Twist1*$^{+/-}$ mouse. The arrowhead denotes that the orientation of the coronal sutures is more 'box-like' compared to wildtype controls. (D) Following injection of 45kDa ovalbumin tracer into the cisterna magna, drainage to the dCLNs is normal in *Twist1*$^{+/-}$ mice that lack coronal suture fusion, whereas *Twist1*$^{+/-}$ mice with coronal suture synostosis have a significant reduction. (D') Quantification of percent area fraction of 45kDa tracer [*Twist1*$^{+/+}$ (n=5); *Twist1*$^{+/-}$ no fusion (n=5); *Twist1*$^{+/-}$ fusion (n=6)]. (E) Following injection of 3kDa Dextran and 45kDa ovalbumin tracer into the cisterna magna, tracer perfusion into the brain is comparable between wildtype controls and *Twist1*$^{+/-}$ mice that lack coronal suture fusion or have very mild partial unilateral fusion, versus *Twist1*$^{+/-}$ mice with near-complete unilateral or bilateral fusion of the coronal sutures. (E') Quantification of percent area fraction of 3kDa Dextran and 45kDa ovalbumin tracer in brain tissue [*Twist1*$^{+/+}$ (n=3); *Twist1*$^{+/-}$ no fusion (n=5); *Twist1*$^{+/-}$ fusion (n=7)]. ns, non-significant, *$p \leq 0.05$,*

**$p\leq0.01$ *One way ANOVA with Tukey's multiple comparison test*. Scale bar: (A and A') 1mm; (C) 5mm; (D) 200μm; (E) 1mm

**Figure S3: *Fgfr2$^{+/S252W}$:Sm22a-Cre* mice have CS, raised intracranial pressure, and hypoplastic dorsal MLVs.** (A) Representative reconstructed computed tomography (CT) scans from two-month-old adults showing normal skull and suture morphology in *Fgfr2$^{+/+}$* control versus *Fgfr2$^{+/S252W}$:Sm22a-Cre* mice with CS. (Middle image) Near-complete and complete fusion of the left and right coronal sutures (arrowheads), respectively, with partial fusion of the sagittal suture (arrowhead). (Right image) Complete fusion of the sagittal suture and bilateral fusion of the coronal sutures. (A') Intracranial pressure is significantly elevated in *Fgfr2$^{+/S252W}$:Sm22a-Cre* mice with CS [*Fgfr2$^{+/+}$* (n=7); *Fgfr2$^{+/S252W}$:Sm22a-Cre* (n=5)]. (B) Representative images showing hypoplasia of dorsal MLVs in *Fgfr2$^{+/S252W}$:Sm22a-Cre* mice with CS. (B') Quantification of percent area fraction Lyve-1 and ovalbumin 45kDa [*Fgfr2$^{+/+}$* (n=5); *Fgfr2$^{+/S252W}$:Sm22a-Cre* (n=9)]. CorS=coronal suture; SS=sagittal suture; TVS=transverse sinus; CoS=confluence of sinuses; SSS=superior sagittal sinus. ***$p\leq0.001$. *two-tailed unpaired t test with Welch's correction.* Scale bar: (A) 5mm; (C) 1mm.

**Figure S4: Homozygous *Twist1$^{+/FLX}$:Sm22a-Cre* mice have raised intracranial pressure and reduced CSF flow to perisinusoidal dura.** (A) Similar to heterozygous *Twist1$^{+/FLX}$:Sm22a-Cre* mice, homozygous *Twist1$^{FLX/FLX}$:Sm22a-Cre* mice with CS have raised intracranial pressure [*Twist1$^{FLX/+}$* (n=7); *Twist1$^{FLX/+}$:Sm22a-Cre* (n=6); *Twist1$^{FLX/FLX}$:Sm22a-Cre* (n=5)]. (B) Representative image showing lack of ovalbumin 45kDa tracer in perisinusoidal dura surrounding MLVs in a 2-month-old *Twist1$^{FLX/FLX}$:Sm22a-Cre* mouse. Quantification of percent area fraction of ovalbumin 45kDa tracer in perisinusoidal dura [*Twist1$^{FLX/+}$* and *Twist1$^{FLX/FLX}$:Sm22a-Cre* (n=4)]. TVS=transverse sinus; SgS=sigmoid sinus; PSS=petrosquamosal sinus. *$p\leq0.05$, **$p\leq0.01$, ***$p\leq0.001$ *One way ANOVA with Tukey's multiple comparison test* (A) and *two-tailed unpaired t test* (B). Data points in A reflecting *Twist1$^{FLX/+}$* and *Twist1$^{FLX/+}$:Sm22a-Cre* mice are from Figure 1. Scale bar=1mm

**Figure S5: *Fgfr2$^{+/S252W}$:Sm22a-Cre* mice with CS have altered CSF flow with reduced perfusion of CSF macromolecules into the brain.** (A) Representative images taken during transcranial live imaging of a 45kDa ovalbumin tracer injected into the cisterna magna of young adult mice. Compared to unaffected littermates (top panel), *Fgfr2$^{+/S252W}$:Sm22a-Cre* mice (bottom panel) show perturbations to CSF tracer flow along preferred dorsal pathways, similar to what is observed in *Twist1$^{+/-}$* mice. (B) Representative images show that perfusion of CSF macromolecules into the brain is reduced in *Fgfr2$^{+/S252W}$:Sm22a-Cre* mice [*Fgfr2$^{+/+}$* (n=5); *Fgfr2$^{+/S252W}$:Sm22a-Cre* (n=6)]. *$p\leq0.01$ *two-tailed unpaired t test*. Scale bar: (A) 5mm; (B) 500μm

**Figure S6: AQP4 is reduced at glial endfeet that wrap around large-caliber vessels.** (A) Coronal sections through the dorsolateral cortex in P17 juveniles. The coverage of AQP4 protein along large-caliber blood vessels labeled with GS-lectin IB4 is reduced in *Twist1*$^{+/-}$ mice (arrowheads). (A') Quantification of percent colocalization of AQP4 protein and lectin [*Twist1*$^{+/+}$ (n=7); *Twist1*$^{+/-}$ (n=6)]. (B) Protein fractions from P17 juvenile mice containing glial endfeet tethered to blood vessels show a reduction in AQP4 protein. (B') Quantification of AQP4 in the vascular fraction (normalized to actin) from three *Twist*$^{+/+}$ and three *Twist1*$^{+/-}$ brains [*Twist1*$^{+/+}$ and *Twist1*$^{+/-}$ (n=3)]. (C) In young *Twist1*$^{+/-}$ adult mice with CS, AQP4 coverage is still reduced along large-caliber vessels (arrowheads). (C') Quantification of percent colocalization of AQP4 protein and lectin [*Twist1*$^{+/+}$ (n=5); *Twist1*$^{+/-}$ (n=6)]. (D) Quantification of percent colocalization of AQP4 protein and lectin in P30 mice treated with saline vehicles or Yoda1. *Twist1*$^{+/-}$*:Yoda1* mice still show a significant reduction of AQP4 protein along large-caliber blood vessels labeled with GS-lectin IB4 compared with controls receiving Yoda1 or saline vehicle [*Twist1*$^{+/+}$ vehicle (n=5); *Twist1*$^{+/+}$ Yoda1 (n=6); *Twist1*$^{+/-}$ vehicle (n=4); *Twist1*$^{+/-}$ Yoda1 (n=5)]. (A', B', C') ***p≤0.01 two-tailed unpaired t test*. (D) ***p≤0.01 one-way ANOVA with Dunnett's multiple comparison test*. Scale bar: (A and C) 200μm

**Figure S7: *Twist1*$^{+/-}$ CS mice do not show changes to astro- or microgliosis.** (A) Representative coronal sections through the cortex and hippocampus stained with GFAP and IBA1. No changes to astro- or microgliosis are detected in *Twist1*$^{+/-}$ craniosynostosis mice, as measured by the percent area fraction of GFAP and IBA1 staining compared to control littermates [*Twist1*$^{+/+}$ (n=5); *Twist1*$^{+/-}$ (n=4)]. (B) Representative images of microglia in the cortex at higher magnification. No significant changes to morphology are detected [*Twist1*$^{+/+}$ (n=3); *Twist1*$^{+/-}$ (n=3)]. *two-tailed unpaired t test with Welch's correction*. Scale bar: (A) 50μm; (B) 100μm

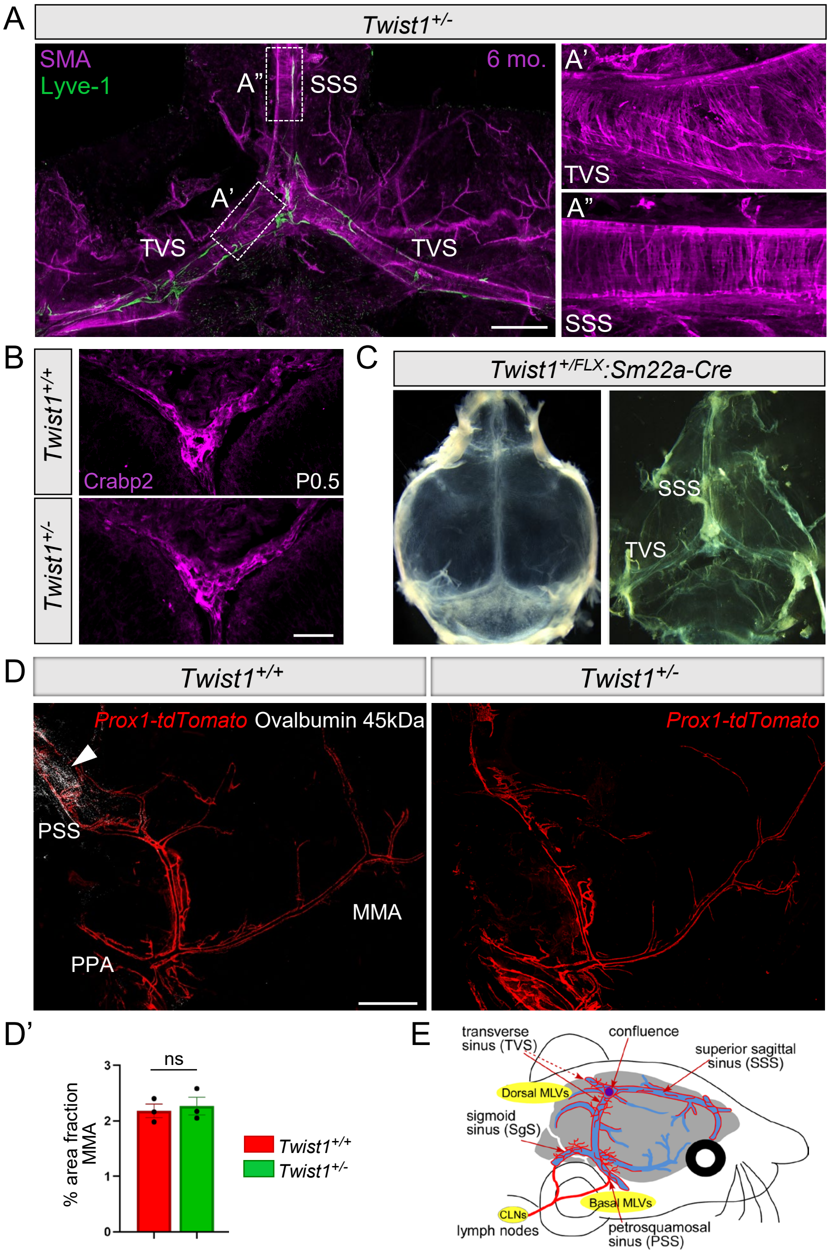


**Figure S1: Dura mater and the venous sinuses are unaffected in *Twist1*$^{+/FLX}$*:Sm22a-Cre* and *Twist1*$^{+/-}$ CS models, and MLVs along the PPA and MMA do not have direct access to CS and develop normally.** (A) Dural scraping from a six-month-old *Twist1*$^{+/-}$ CS mouse (n=3) with regressed dorsal MLVs. The transverse sinuses are both present. Smooth muscle coverage is normal on the transverse (A') and superior sagittal sinuses (A"). (B) Coronal cross section at the dorsal midline of a P0.5 mouse labeled with Crabp2, which marks dura and the underlying arachnoid. The dura and arachnoid membranes show normal Crabp2 expression and are not hypoplastic in *Twist1*$^{+/-}$ CS mice. (C) Dorsal craniotomy and dural flat mount. The dura is easily peeled off the skull in *Twist*$^{+/-}$ and *Twist1*$^{+/FLX}$*:Sm22a-Cre* mice (shown). Representative images from young adult mice aged 2-4 months. (D and D') The percent area fraction of *Prox1-tdTomato* signal along the MMA is normal in *Twist1*$^{+/-}$ CS mice compared with unaffected littermates [*Twist1*$^{+/+}$ (n=3); *Twist1*$^{+/-}$ (n=3)]. Note that tracer is present (arrowhead) in perisinusoidal dura enveloping the PSS (petrosquamosal sinus) but is absent in dura surrounding the PPA (pterygopalatine artery) and MMA (middle meningeal artery). (E) Schematic overview of dorsal and basal perisinusoidal MLVs in mouse. ns, non-significant, *two-tailed unpaired t test.* Scale bar: (A) 250mm; (B) 50mm; (D) 1mm.

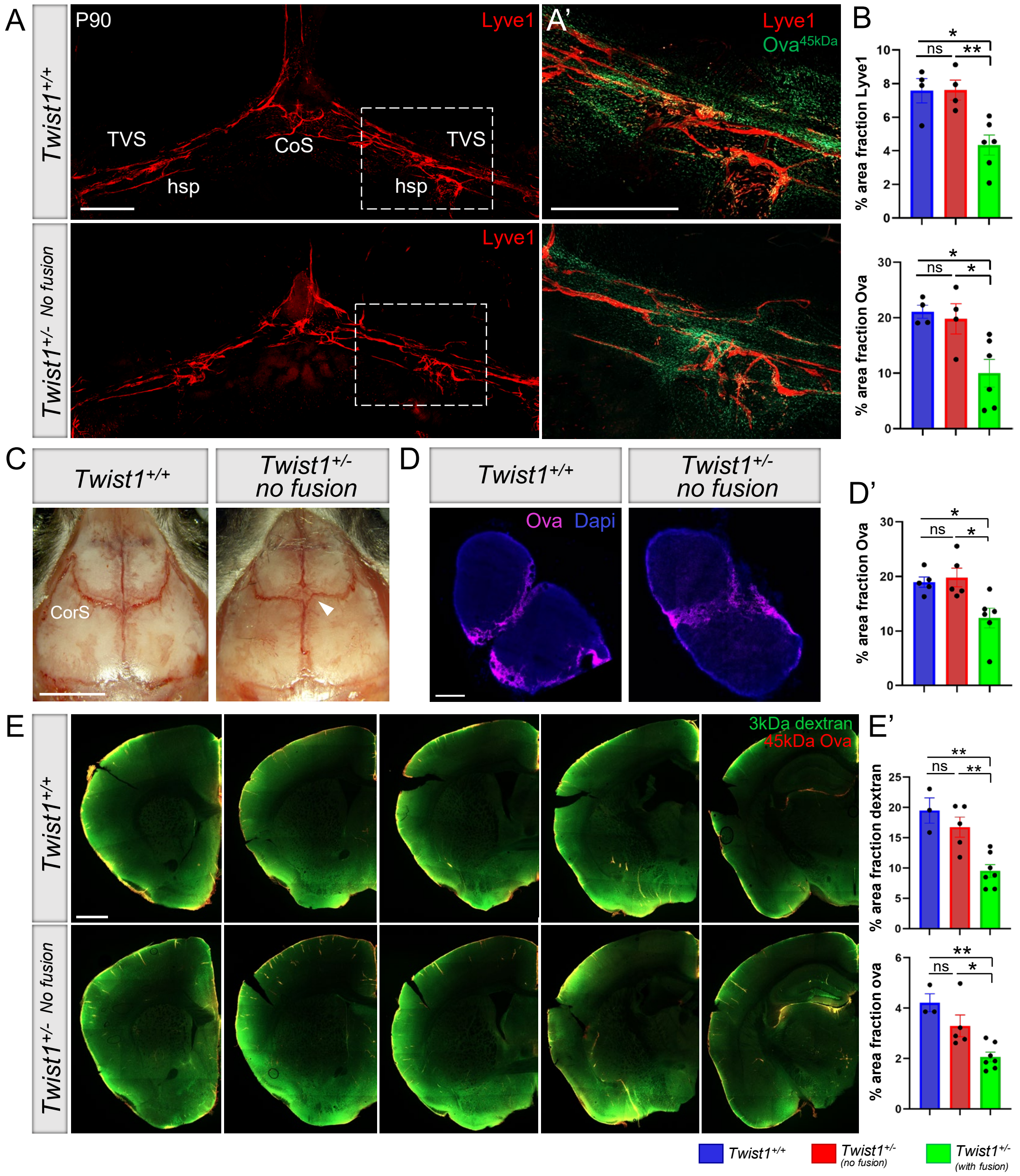


**Figure S2: MLV networks, flow to perisinusoidal dura, drainage to the dCLNs, and brain-CSF perfusion are unaffected in *Twist1$^{+/-}$* mice without suture fusion.** (A) Representative images show that dorsal MLV networks, stained with Lyve-1, and hotspots along the transverse sinus (TVS) develop normally in *Twist1$^{+/-}$* mice that lack coronal suture fusion. (A') Following injection of 45kDa ovalbumin tracer into the cisterna magna, tracer deposition in perisinusoidal dura surrounding MLVs and hotspots is comparable between controls and *Twist1$^{+/-}$* mice that lack fusion of the coronal sutures, versus *Twist1$^{+/-}$* mice that have coronal suture synostosis. (B) Quantification of percent area fraction Lyve-1 and ovalbumin 45kDa [*Twist1$^{+/+}$* (n=4); *Twist1$^{+/-}$* no fusion (n=5); *Twist1$^{+/-}$* fusion (n=6)]. (C) Skull overviews showing patent coronal sutures (CorS) in a *Twist1$^{+/-}$* mouse. The arrowhead denotes that the orientation of the coronal sutures is more 'box-like' compared to wildtype controls. (D) Following injection of 45kDa ovalbumin tracer into the cisterna magna, drainage to the dCLNs is normal in *Twist1$^{+/-}$* mice that lack coronal suture fusion, whereas *Twist1$^{+/-}$* mice with coronal suture synostosis have a significant reduction. (D') Quantification of percent area fraction of 45kDa tracer [*Twist1$^{+/+}$* (n=5); *Twist1$^{+/-}$* no fusion (n=5); *Twist1$^{+/-}$* fusion (n=6)]. (E) Following injection of 3kDa Dextran and 45kDa ovalbumin tracer into the cisterna magna, tracer perfusion into the brain is comparable between wildtype controls and *Twist1$^{+/-}$* mice that lack coronal suture fusion or have very mild partial unilateral fusion, versus *Twist1$^{+/-}$* mice with near-complete unilateral or bilateral fusion of the coronal sutures. (E') Quantification of percent area fraction of 3kDa Dextran and 45kDa ovalbumin tracer in brain tissue [*Twist1$^{+/+}$* (n=3); *Twist1$^{+/-}$* no fusion (n=5); *Twist1$^{+/-}$* fusion (n=7)]. ns, non-significant, *$*p\leq0.05$, $**p\leq0.01$ One way ANOVA with Tukey's multiple comparison test*. Scale bar: (A and A') 1mm; (C) 5mm; (D) 200µm; (E) 1mm

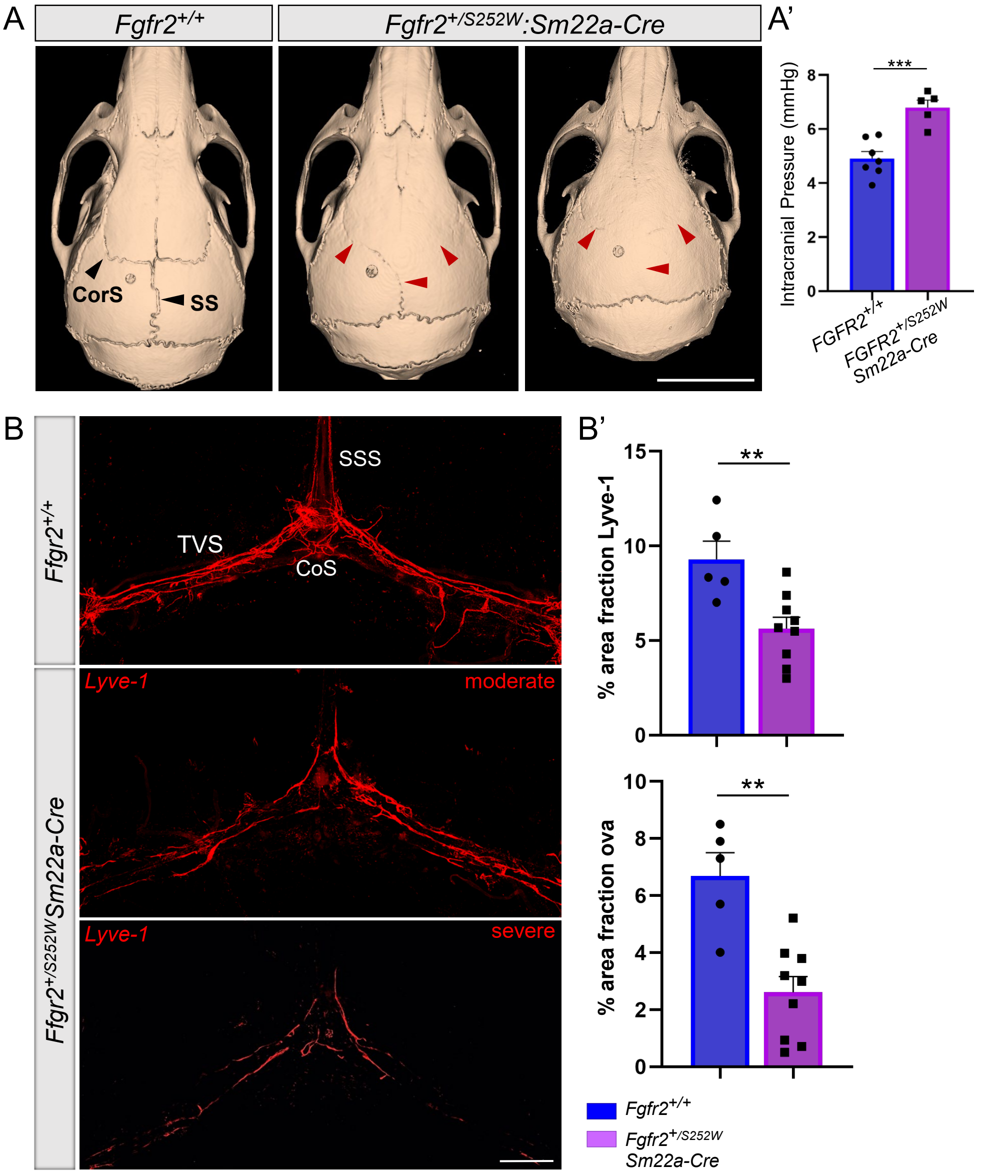


**Figure S3: *Fgfr2+/S252W:Sm22a-Cre* mice have CS, raised intracranial pressure, and hypoplastic dorsal MLVs.** (A) Representative reconstructed computed tomography (CT) scans from two-month-old adults showing normal skull and suture morphology in *Fgfr2+/+* control versus *Fgfr2+/S252W:Sm22a-Cre* mice with CS. (Middle image) Near-complete and complete fusion of the left and right coronal sutures (arrowheads), respectively, with partial fusion of the sagittal suture (arrowhead). (Right image) Complete fusion of the sagittal suture and bilateral fusion of the coronal sutures. (A') Intracranial pressure is significantly elevated in *Fgfr2+/S252W:Sm22a-Cre* mice with CS [*Fgfr2+/+* (n=7); *Fgfr2+/S252W:Sm22a-Cre* (n=5)]. (B) Representative images showing hypoplasia of dorsal MLVs in *Fgfr2+/S252W:Sm22a-Cre* mice with CS. (B') Quantification of percent area fraction Lyve-1 and ovalbumin 45kDa [*Fgfr2+/+* (n=5); *Fgfr2+/S252W:Sm22a-Cre* (n=9)]. CorS=coronal suture; SS=sagittal suture; TVS=transverse sinus; CoS=confluence of sinuses; SSS=superior sagittal sinus. ****p≤0.001. two-tailed unpaired t test with Welch's correction.* Scale bar: (A) 5mm; (C) 1mm.

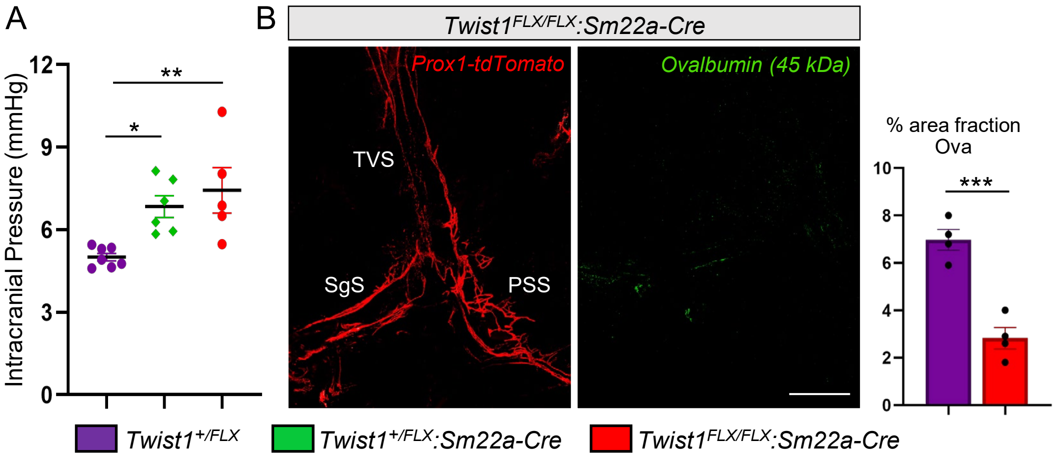


**Figure S4: Homozygous *Twist1*$^{+/FLX}$*:Sm22a-Cre* mice have raised intracranial pressure and reduced CSF flow to perisinusoidal dura.** (A) Similar to heterozygous *Twist1*$^{+/FLX}$*:Sm22a-Cre* mice, homozygous *Twist1*$^{FLX/FLX}$*:Sm22a-Cre* mice with CS have raised intracranial pressure [*Twist1*$^{FLX/+}$ (n=7); *Twist1*$^{FLX/+}$*:Sm22a-Cre* (n=6); *Twist1*$^{FLX/FLX}$*:Sm22a-Cre* (n=5)]. (B) Representative image showing lack of ovalbumin 45kDa tracer in perisinusoidal dura surrounding MLVs in a 2-month-old *Twist1*$^{FLX/FLX}$*:Sm22a-Cre* mouse. Quantification of percent area fraction of ovalbumin 45kDa tracer in perisinusoidal dura [*Twist1*$^{FLX/+}$ and *Twist1*$^{FLX/FLX}$*:Sm22a-Cre* (n=4)]. TVS=transverse sinus; SgS=sigmoid sinus; PSS=petrosquamosal sinus. *$p \leq 0.05$, **$p \leq 0.01$, ***$p \leq 0.001$ *One way ANOVA with Tukey's multiple comparison test* (A) and *two-tailed unpaired t test* (B). Data points in A reflecting *Twist1*$^{FLX/+}$ and *Twist1*$^{FLX/+}$*:Sm22a-Cre* mice are from Figure 1. Scale bar=1mm

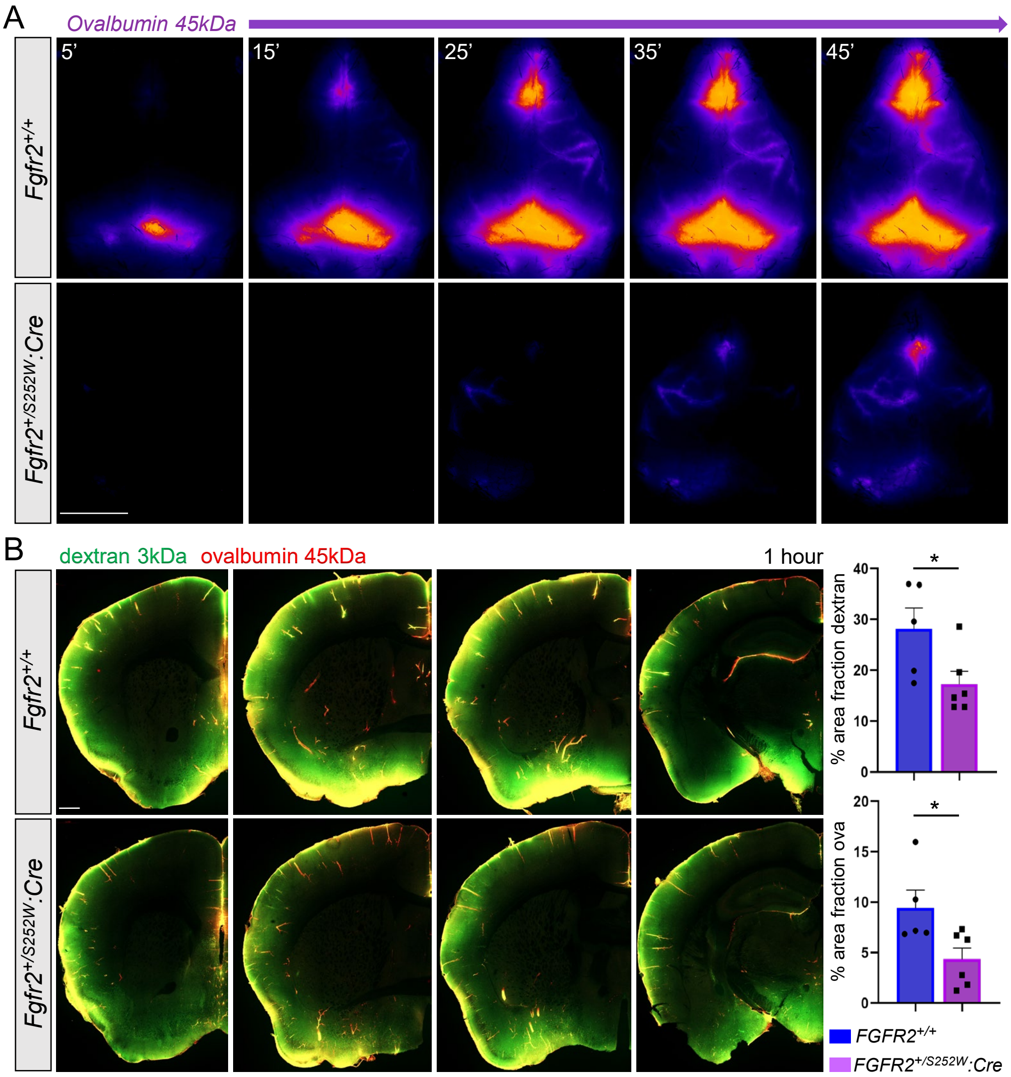


**Figure S5: *Fgfr2$^{+/S252W}$:Sm22a-Cre* mice with CS have altered CSF flow with reduced perfusion of CSF macromolecules into the brain.** (A) Representative images taken during transcranial live imaging of a 45kDa ovalbumin tracer injected into the cisterna magna of young adult mice. Compared to unaffected littermates (top panel), *Fgfr2$^{+/S252W}$:Sm22a-Cre* mice (bottom panel) show perturbations to CSF tracer flow along preferred dorsal pathways, similar to what is observed in *Twist1$^{+/-}$* mice. (B) Representative images show that perfusion of CSF macromolecules into the brain is reduced in *Fgfr2$^{+/S252W}$:Sm22a-Cre* mice [*Fgfr2$^{+/+}$* (n=5); *Fgfr2$^{+/S252W}$:Sm22a-Cre* (n=6)]. *$p \leq 0.01$ *two-tailed unpaired t test*. Scale bar: (A) 5mm; (B) 500µm

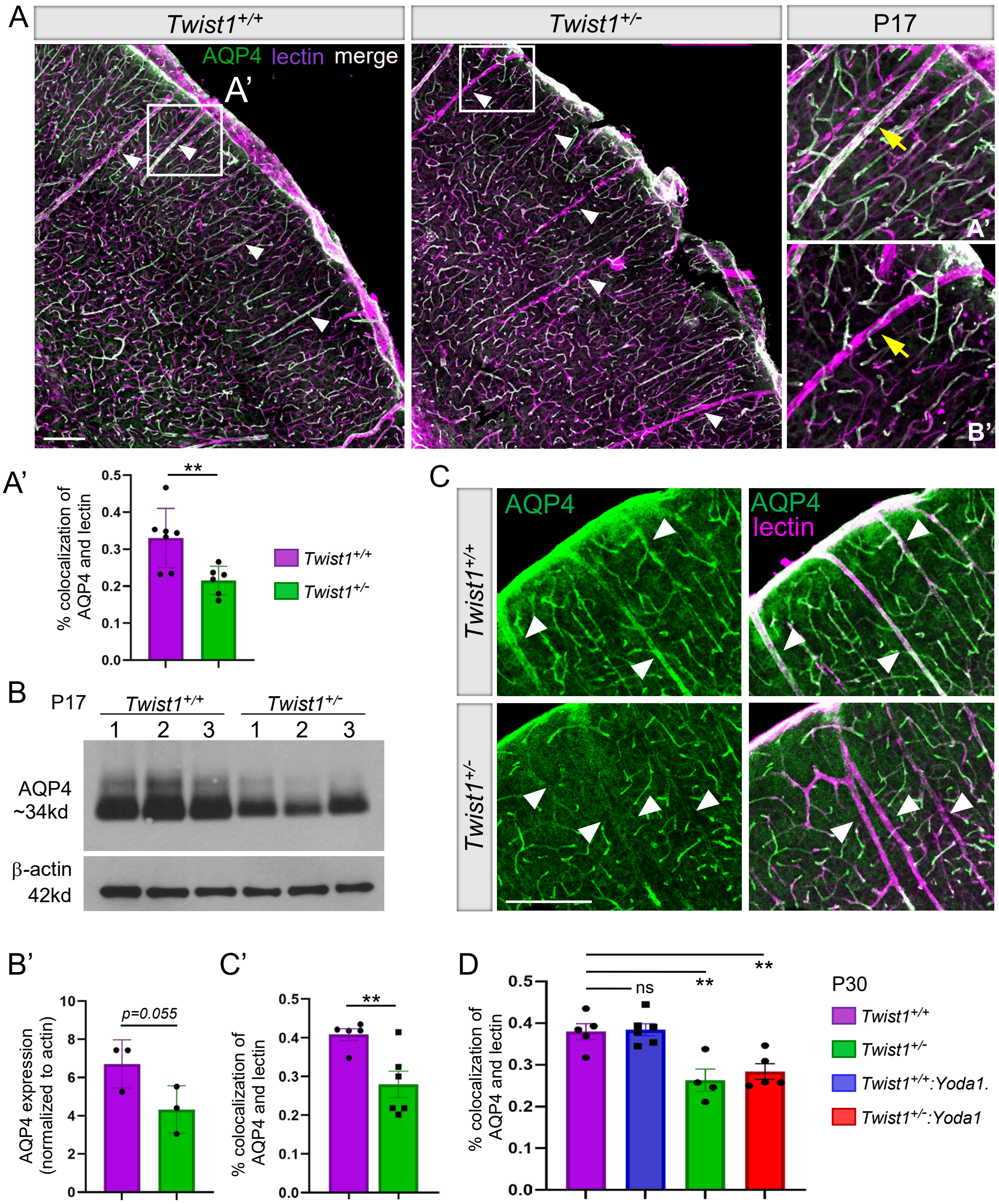


**Figure S6: AQP4 is reduced at glial endfeet that wrap around large-caliber vessels.** (A) Coronal sections through the dorsolateral cortex in P17 juveniles. The coverage of AQP4 protein along large-caliber blood vessels labeled with GS-lectin IB4 is reduced in *Twist1*$^{+/-}$ mice (arrowheads). (A') Quantification of percent colocalization of AQP4 protein and lectin [*Twist1*$^{+/+}$ (n=7); *Twist1*$^{+/-}$ (n=6)]. (B) Protein fractions from P17 juvenile mice containing glial endfeet tethered to blood vessels show a reduction in AQP4 protein. (B') Quantification of AQP4 in the vascular fraction (normalized to actin) from three *Twist*$^{+/+}$ and three *Twist1*$^{+/-}$ brains [*Twist1*$^{+/+}$ and *Twist1*$^{+/-}$ (n=3)]. (C) In young *Twist1*$^{+/-}$ adult mice with CS, AQP4 coverage is still reduced along large-caliber vessels (arrowheads). (C') Quantification of percent colocalization of AQP4 protein and lectin [*Twist1*$^{+/+}$ (n=5); *Twist1*$^{+/-}$ (n=6)]. (D) Quantification of percent colocalization of AQP4 protein and lectin in P30 mice treated with saline vehicles or Yoda1. *Twist1*$^{+/-}$*:Yoda1* mice still show a significant reduction of AQP4 protein along large-caliber blood vessels labeled with GS-lectin IB4 compared with controls receiving Yoda1 or saline vehicle [*Twist1*$^{+/+}$ vehicle (n=5); *Twist1*$^{+/+}$ Yoda1 (n=6); *Twist1*$^{+/-}$ vehicle (n=4); *Twist1*$^{+/-}$ Yoda1 (n=5)]. (A', B', C') ***$p \leq 0.01$ two-tailed unpaired t test*. (D) ***$p \leq 0.01$ one-way ANOVA with Dunnett's multiple comparison test*. Scale bar: (A and C) 200μm

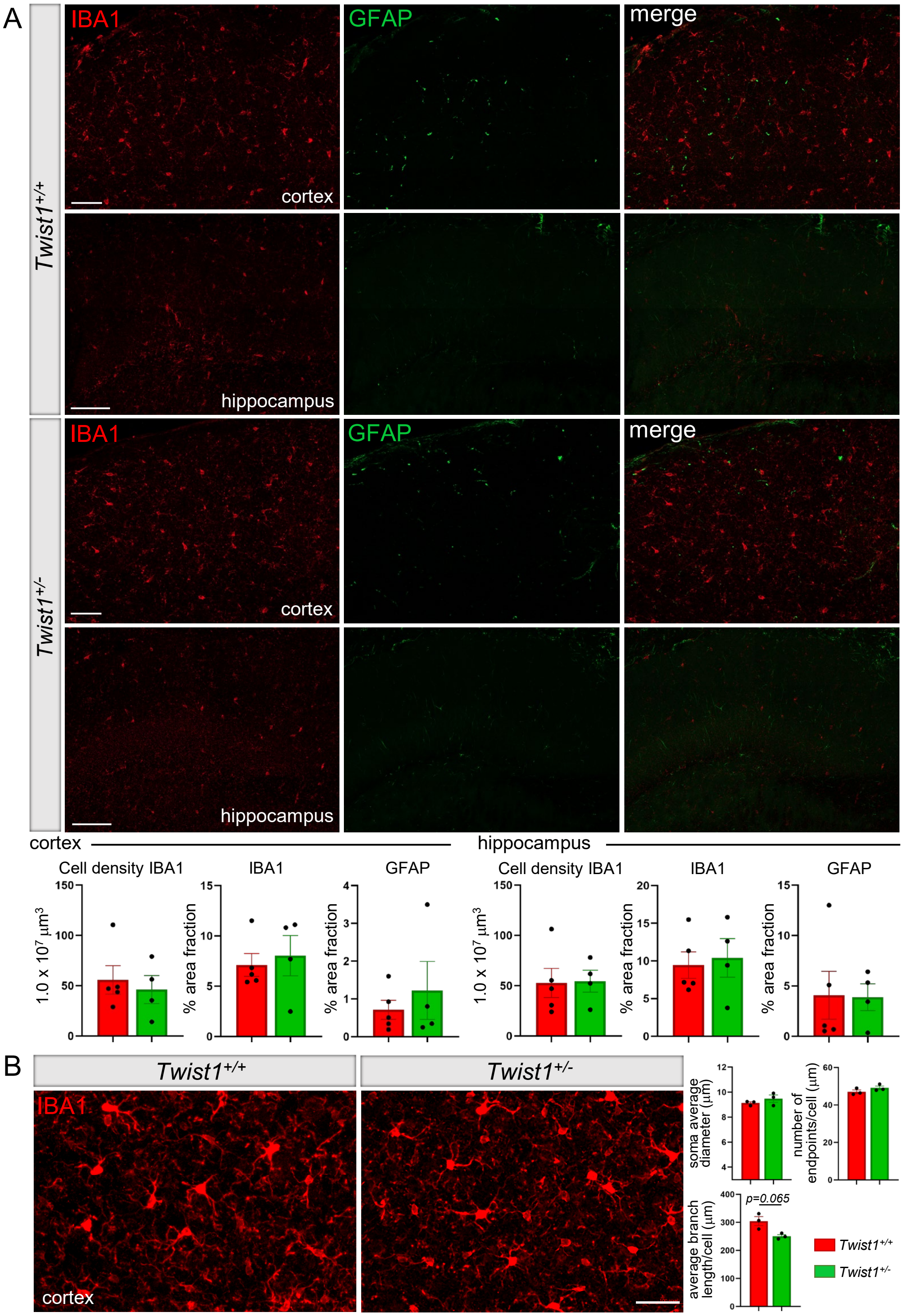


**Figure S7:** ***Twist1**$^{+/-}$* **CS mice do not show changes to astro- or microgliosis.** (A) Representative coronal sections through the cortex and hippocampus stained with GFAP and IBA1. No changes to astro- or microgliosis are detected in *Twist1*$^{+/-}$ craniosynostosis mice, as measured by the percent area fraction of GFAP and IBA1 staining compared to control littermates [*Twist1*$^{+/+}$ (n=5); *Twist1*$^{+/-}$ (n=4)]. (B) Representative images of microglia in the cortex at higher magnification. No significant changes to morphology are detected [*Twist1*$^{+/+}$ (n=3); *Twist1*$^{+/-}$ (n=3)]. *two-tailed unpaired t test with Welch's correction.* Scale bar: (A) 50mm; (B) 100mm
